# The crystal structure of the activin B:Fst288 complex and computational insights into the broad antagonistic activity and specificity of follistatin

**DOI:** 10.1101/2025.11.14.688564

**Authors:** Lucija Hok, Ryan G. Walker, James A. Howard, Eleanor M. Mast, Chandramohan Kattamuri, Erich J. Goebel, Thomas B. Thompson

## Abstract

Members of the TGF-β family regulate diverse biological processes, and their activity is tightly controlled by extracellular antagonists such as follistatin 288 (Fst288). Fst288 potently inhibits activins A and B, GDF8, and GDF11. Among these, ActB is the least structurally and functionally characterized, which limits a full understanding of Fst288’s broad activity and its specificity for this TGF-β subgroup. To address this, we solved the crystal structure of the ActB:Fst288 complex. As in other complexes, two Fst288 molecules surround ActB, occluding both receptor-binding sites. However, ActB engages Fst288 differently: its fingers bind most strongly the ND domain, while its acidic fingertips uniquely contact FSD3 disrupting interactions between the two Fst288 molecules and reducing the cooperativity seen in other complexes. Computational analysis revealed that, although ActB exhibits higher interaction enthalpy, entropic penalty lowers its overall affinity compared to ActA, consistent with experimentally measured *K*_d_ values. We therefore propose that broad ligand inhibition by Fst288 arises from variations in binding interactions and the utilization of cooperativity when direct contacts are insufficient for high affinity. In cellular contexts, antagonistic effectiveness is further impacted by interactions between complexes and the extracellular matrix, resulting in comparable *in vitro* IC_50_ values. Finally, sequence comparisons with non-binding TGF-βs indicate that Fst288 specificity originates from ionic contacts with acidic ligand fingertips which likely initiate recognition, while interactions at the type I and II interfaces stabilize the complex. This study provides deeper insight into Fst288’s regulation of activin signaling and paves the way for designing inhibitors with desired selectivity.

## Introduction

The transforming growth factor-β (TGF-β) family of cytokines regulates many physiological processes such as cell proliferation, differentiation, and homeostasis. Members of this family are broadly categorized into three subgroups: TGF-βs, activins/inhibins, and bone morphogenetic proteins (BMPs). TGF-β ligands are typically disulfide-linked dimers with a characteristic cysteine-knot fold that signal through the canonical pathway by forming ternary complexes with two type II and two type I receptors. Type II receptors are constitutively active serine/threonine kinases that phosphorylate type I receptors upon ligand binding. Activated type I receptors phosphorylate intracellular Smad proteins which then translocate into the nucleus and regulate gene expression (1).

The actions of TGF-β ligands are tightly regulated by extracellular antagonists with diverse specificity profiles – some inhibit only a narrow range of ligands, while others exhibit broader activity – therefore showing distinct biological functions. Among the latter is follistatin 288 (Fst288), a multidomain glycoprotein composed of an N-terminal domain (ND) and three follistatin domains (FSD1–3), which are further subdivided into EGF-like and Kazal protease inhibitor-like domains (Fig. 1A and S1). Together with Fst315 and Fstl3, it belongs to the group of Fst domain-containing antagonists that differ in the architecture, binding characteristics and specificity for ligands (2, 3). More specifically, Fst315 is a Fst288 isoform with an acidic C-terminal tail, while Fstl3 has a domain structure similar to that of Fst288, but lacks FSD3.

**Figure 1.**
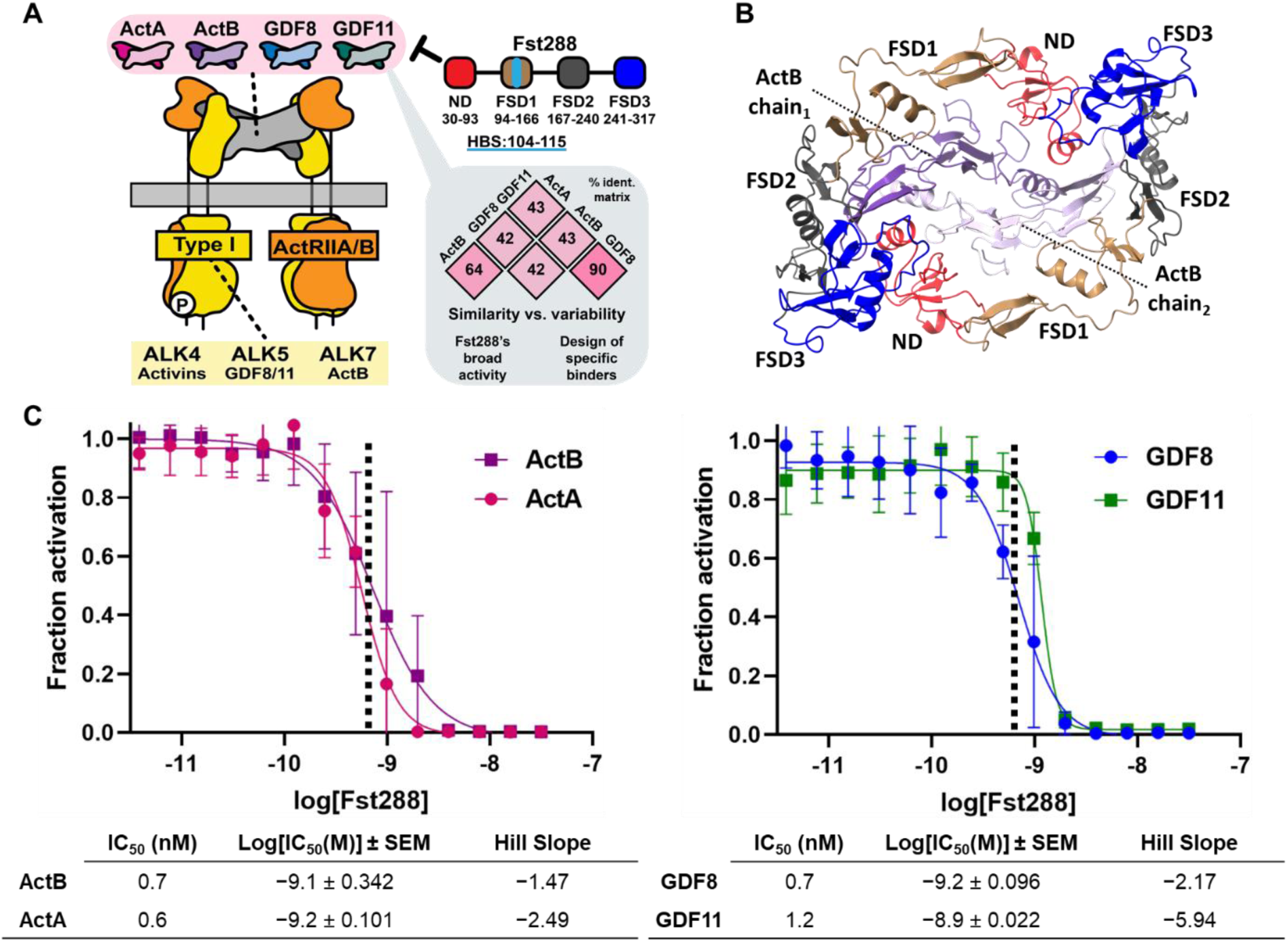
Mechanism and Inhibitory Activity of Fst288 Toward Activins. A) Schematic of activins signaling inhibition by Fst288 and sequence identity matrix (%) calculated for the growth factor domain. B) Top view of the crystal structure of the ActB:Fst288 complex. C) Luciferase reporter assays showing inhibitory activity following titration of Fst288 against a constant concentration (0.62 nM, dashed line) of activin B (purple) and activin A (pink) of the left, and GDF8 (blue), GDF11 (green) on the right, in HEK293 (CAGA)_12_ cells.

The widespread defects in the *Fst* knockout mice, such as impaired breathing and developmental malformations in the whiskers, teeth and muscles, affirm Fst288’s broad antagonistic role (4). Initially identified as a strong inhibitor of activin A (ActA), Fst288 also inhibits activin B (ActB), GDF8, and GDF11 at nanomolar concentrations (5, 6). It has limited impact on BMP signaling and does not inhibit the TGF-β subgroup (7). Crystal structures of Fst-type molecules in complex with ActA, GDF8, or GDF11 show that neutralization involves two Fst molecules blocking both receptor-binding sites (5, 8–12).

Numerous studies have investigated the importance of individual Fst domains to activins’ antagonism through mutational and domain-swap studies (10, 13–20). Both ND and FSD1–2 are required for ActA inhibition (13), while an ND-FSD1-FSD1 construct specifically antagonizes GDF8 (14, 16, 19). Point mutations in the ND and FSD2 were shown equally detrimental to antagonism, suggesting that affinity for ligands is spread out over both domains (14, 20). FSD3, although not directly contacting the ligand, enhances binding and can rescue antagonism in mutated forms of Fstl3 and Fst288 (10, 13, 14). Moreover, FSD3 confers cooperativity by interacting with the ND of the adjacent Fst288 molecule in a head-to-tail configuration. The lack of cooperativity in Fstl3 and ΔFSD3 constructs lead to the speculation that the ND–FSD3 interaction underlies Fst288’s broad activity since it is ligand independent (14). However, differential antagonistic potential towards different TGF-β ligands indicates that ligands’ specific structural features are equally important in defining the final sensitivity.

While, among activins, ActA, GDF8, and GDF11 are structurally and functionally well characterized in both their apo forms and in complexes with various binding partners (5, 8–12, 15, 21–27), much less is known about ActB or its interactions with antagonists, as available structural information is limited to the ActB:BMPR2 complex (23). To address this gap and better understand Fst288’s antagonistic activity, we determined the crystal structure of ActB in complex with Fst288 (PDB ID: 9Z3T). This allowed us to make a comparative analysis with previously resolved activin:Fst structures and show that ActB engages Fst288 differently at the type I receptor binding site, relying more on its fingers than other activins, resulting in a higher interaction energy estimated by computational methods. Moreover, ActB directly interacts with FSD3 through its highly acidic fingertips, disrupting the ND–FSD3 interaction and disabling cooperativity. This indicates Fst288 can adapt its binding mode dependent on the ligand: when strong contacts with individual domains are missing, Fst288 adjusts its conformation and compensates by utilizing cooperativity through the ND–FSD3 interaction. Furthermore, a broader sequence comparison with BMPs and TGF-βs lead us to propose a model in which the acidic fingertips drive an initial recognition, while the FSD2 and ND binding stabilizes the interaction and fine-tune affinity. The structural characterization of the last remaining high-affinity activin:Fst288 complex provides new insights into the regulation of TGF-β signaling by endogenous antagonists, and offers a structural basis for designing inhibitors with desired selectivity to treat pathophysiological conditions associated with elevated ligand levels (28–32) (Fig. 1A).

## Results

### The structure of the ActB:Fst288 complex

The complex of the ActB dimer bound to two Fst288 molecules was resolved using X-ray crystallography to 2.7 Å (Table 1) and represents the first structure of ActB in complex with a known antagonist. Similar to previously published activin:Fst structures (5, 8–12), two molecules of Fst288 surround the dimeric ligand burying both type I and type II receptor binding sites (Fig. 1B). The ND domain occupies type I receptor site contacting concave region of the ligand fingers and the wrist helix of the opposite chain, while the FSD2 and, to a lesser extent, the FSD1 domains bind to the convex surface of the fingers overlapping with the type II binding epitope.

**Table 1.**
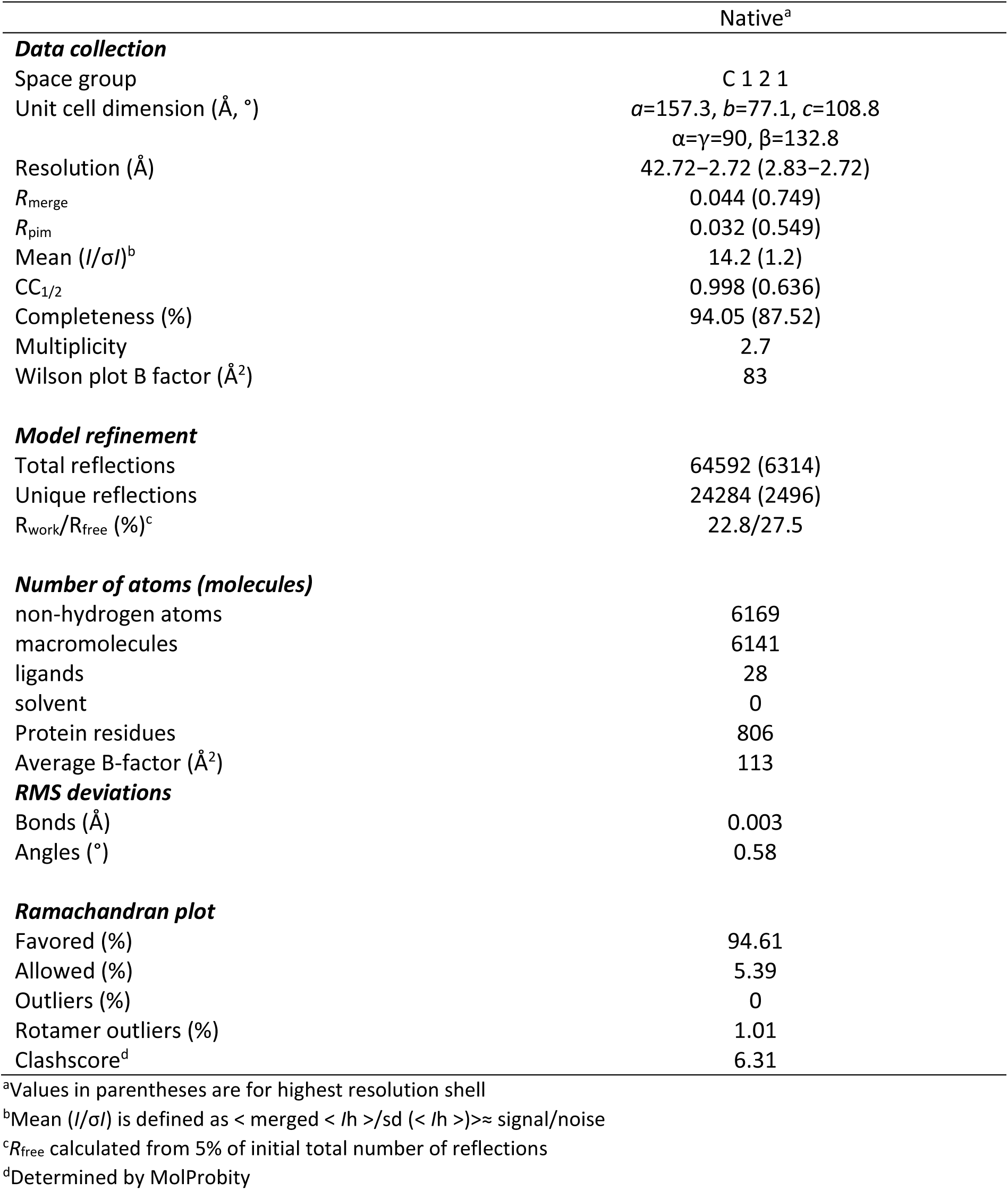
Data Collection and Refinements statistics (Molecular Replacement).

|  | Native<sup>a</sup> |
| --- | --- |
| **Data collection** |  |
| Space group | C 1 2 1 |
| Unit cell dimension (Å, °) | $a=157.3$, $b=77.1$, $c=108.8$<br>$\alpha=\gamma=90$, $\beta=132.8$ |
| Resolution (Å) | 42.72–2.72 (2.83–2.72) |
| $R_{\text{merge}}$ | 0.044 (0.749) |
| $R_{\text{pim}}$ | 0.032 (0.549) |
| Mean ($I/\sigma I$)<sup>b</sup> | 14.2 (1.2) |
| $CC_{1/2}$ | 0.998 (0.636) |
| Completeness (%) | 94.05 (87.52) |
| Multiplicity | 2.7 |
| Wilson plot B factor (Å<sup>2</sup>) | 83 |
| **Model refinement** |  |
| Total reflections | 64592 (6314) |
| Unique reflections | 24284 (2496) |
| $R_{\text{work}}/R_{\text{free}}$ (%)<sup>c</sup> | 22.8/27.5 |
| **Number of atoms (molecules)** |  |
| non-hydrogen atoms | 6169 |
| macromolecules | 6141 |
| ligands | 28 |
| solvent | 0 |
| Protein residues | 806 |
| Average B-factor (Å<sup>2</sup>) | 113 |
| **RMS deviations** |  |
| Bonds (Å) | 0.003 |
| Angles (°) | 0.58 |
| **Ramachandran plot** |  |
| Favored (%) | 94.61 |
| Allowed (%) | 5.39 |
| Outliers (%) | 0 |
| Rotamer outliers (%) | 1.01 |
| Clashscore<sup>d</sup> | 6.31 |

<sup>a</sup>Values in parentheses are for highest resolution shell<sup>b</sup>Mean ( $I/\sigma I$ ) is defined as $\langle \text{merged} \langle I/h \rangle / \text{sd} \langle I/h \rangle \rangle \approx \text{signal/noise}$ <sup>c</sup> $R_{\text{free}}$ calculated from 5% of initial total number of reflections<sup>d</sup>Determined by MolProbity

### Fst288 strongly inhibits activin ligands

Since Fst288 binds activin ligands in a similar manner, blocking both receptor epitopes, before delving deeper into the structural peculiarities of the binding, we first wanted to determine whether Fst288 differs in its effectiveness at inhibiting ActB, ActA, GDF8, and GDF11. For that purpose, we used a HEK293 (CAGA)_12_ luciferase assay to measure inhibitory activity by titrating the antagonist against a constant concentration of each ligand. The results show that Fst288 similarly inhibits all studied ligands at nanomolar concentrations (Fig. 1C).

### Unique features of the ActB:Fst288 interaction

ActB consists of two inhibin β_B_ chains whose growth factor domains share 64% sequence identity with the β_A_ that compose ActA, (33) while GDF8 and GDF11 are ∼40% identical to the β chains, but are closely related to each other, sharing 90% sequence identity (12) (Fig. 1A). Despite higher similarity to ActA, the Fst288 binding of ActB more closely resembles that of GDF8 and GDF11, forming more contacts and burying an additional 635 Å² of surface area (Fig. 2A). However, in several respects, the Fst288 interaction with ActB is unique. One notable difference is the distribution of contacts between the ND and the two ligand chains. While in other complexes the ND primarily interacts with the prehelix loop and wrist helix of chain_2_, in ActB the ND engages both chains equally, with the largest contact area formed between the ND helix and the fingers of chain_1_ (Fig. 2A). Even more strikingly, unlike in other activin complexes where FSD3 either does not or interacts only minimally with the ligand, in the ActB complex FSD3 directly contacts ligand’s fingertips. Interestingly, the type II receptor binding epitope is more similar among complexes than the type I. However, in ActA, FSD1 is vertically displaced downward toward fingers 1 and 2, resulting in a somewhat smaller contact area (Fig. S2).

**Figure 2.**
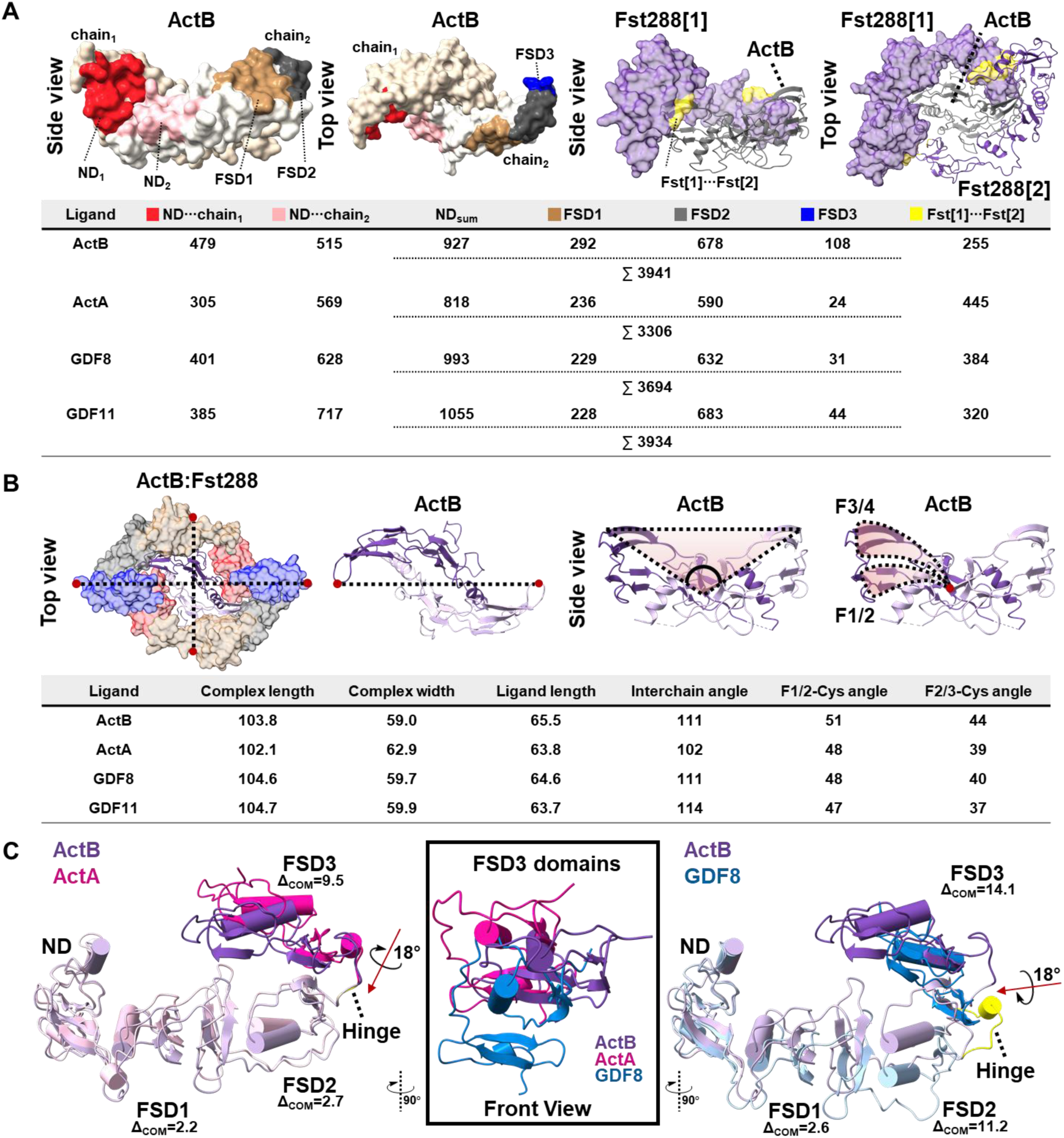
Structural comparison of activin:Fst288 complexes. A) Buried surface area (in Å²) between ActB and individual Fst domains: ND with chain_1_ in red, ND with chain_2_ in pink, FSD1 in brown, FSD2 in grey, FSD3 in blue; and between the two Fst in yellow. Calculated values (BSA ≥ 15 Å²) for activins are given in the table below. B) Geometrical parameters for the complexes: overall complex length and width (in Å) measured between K243_1_-K243_2_ and C106_1_-C106_2_, respectively; ligand length (in Å) (E388_1_-E388_2_ in ActB, G407_1_-G407_2_ in ActA, K356_1_-K356_2_ in GDF8, K388_1_-K388_2_ in GDF11); interchain angle (in °) (V392_1_-C372_1_-V392_2_ in ActB, I410_1_-C390_1_-I410_2_ in ActA, I359_1_-C339_1_-I359_2_ in GDF8, I391_1_-C371_1_-I391_2_ in GDF11); the angles (in °) of fingertips 1 (D319 in ActB, D337 in ActA, D296 in GDF8, D328 in GDF11) and 2 (E388 in ActB, G407 in ActA, K356 in GDF8, K388 in GDF11) with the imaginary plane through the central disulfide bridge. C) Dynamics of Fst288 in the ActB complex compared to ActA (left) and GDF8 (right), aligned through the ND. Red arrows indicate hinge axis and Δ_COM_ values (in Å) represent the displacement of the centers of mass of individual domains in ActA and GDF8 relative to ActB. Center: front views of the FSD3 domains from ActB, ActA, and GDF8 (left and right images rotated vertically by 90°).

### The role of ligand and Fst288 flexibility in forming high-affinity complexes

Crystal structures of activin ligands resolved to date have shown that the growth factors can adopt very different interchain conformations due to the wrist helix being only loosely tethered to the fingers of the opposite monomer (5, 8–12, 15, 21–26). However, how this flexibility influences interactions with binding partners remains unknown. In the ActB:Fst288 complex, the ligand has a conformation similar to that of GDF8 and GDF11, with the interchain angle of 111°, while ActA adopts a ∼10° more closed conformation which results in a vertical shift of FSD1 toward fingers 1 and 2 (*vide supra*; Fig. 2B). Despite these differences, other geometrical features, such as fingertip curling, lead to comparable complex length and similar overall shape.

To explore ligand flexibility, we performed molecular dynamics simulations (MDS) on both the Fst288-bound complexes and their apo forms. In the apo MDS, ActA and GDF8 showed slightly greater flexibility than their respective complexes, but their interchain conformations remained unchanged, suggesting that these structures represent stable local minima that persist even in the absence of Fst288 (Fig. S3). In contrast, the ActB apo simulation displayed substantial root mean square deviation (RMSD) fluctuations, indicating pronounced global rearrangements. The interchain angle analysis showed a conformational transition from the closed state seen in the Fst288 complex (∼111°) to a more open state (135°) which resembles the ActB:BMPR2 crystal structure (142°) (23). Although Fst288 binding stabilizes ActB, the prehelix and wrist helix region remains the most dynamic among the ligands, as evidenced by the largest occupied volume during MDS (Fig. S3). It is possible that, in response to wrist region flexibility, the ND domain repositions itself closer to the more stable ActB fingers resulting in the increased contact area with chain_1_. Interestingly, computational swapping the prehelix and wrist helix regions between ActA and ActB altered ligand dynamics: the ActA^B^ chimera became more flexible, while ActB^A^ showed increased stability compared to their wild-type forms (Fig. S4). However, the ActA^B^ mutant did not transition to the open conformation, suggesting that the wrist interaction with fingers of the opposing chain also contributes to the overall shape.

On the other side, Fst288 also features flexibility due to loops connecting its domains. When Fst288 from crystal structures were aligned through the ND, a different positioning, especially in the FSD3 domain, was observed (Fig. 2C). Similar was seen in the MD simulations of the ActB:Fst288 complex, where FSD3 shifts even closer to the ligand, stabilizing ionic interactions between the ActB fingertip 2 and positively charged residues on FSD3 (Fig. S5). MDS of Fst288 without ligands revealed substantial structural dynamics, primarily driven by ionic interactions between positively charged residues on FSD1 and negatively charged residues on FSD3 (Fig. S6). This indicates that Fst288 possesses enough conformational plasticity to accommodate and stabilize a range of activin ligands.

### ND domain interacts more extensively with ActB chain_1_ compared to other activin ligands

Having characterized the global structural differences among the complexes, we next examined the specific local features in greater detail. As expected from the buried surface area, the ND helix is positioned 2.3 Å deeper into the type I site of ActB than in ActA, resulting in more extensive contacts with chain_1_ (Fig. 3A). This is enabled by the absence of steric clashes with the prehelix region on chain_2_, which in ActA is ordered as an α′ helix, but remains disordered in ActB. Consequently, M79 protrudes deeper into the hydrophobic pocket between the prehelix loop and the wrist helix, also allowing the formation of CH···π interactions between ND I80 and the wrist helix (Fig. 3B). This hydrophobic interaction appears to be important for stabilizing a helical secondary structure in the type I receptor site, as Alk4 has V76, Alk5 I78 and Alk7 V73 at the same position (26, 27). Interestingly, in the ActA complex with Fstl3, where the α′ helix is not formed, the ND helix extends deeper into the concave pocket as in the ActB:Fst288 (10). A similar ND helix position is also adopted in the GDF8 and GDF11 complexes with Fst288, likely due to a shorter prehelix loop (Fig. S7A). However, certain differences, such as the absence of a hydrogen bond analogous to ActB[R394]···ND[F81], due to a glycine at the equivalent position, result in a slightly distant helix position. At the ligand fingertips, structural peculiarities become even more pronounced. In ActB, D387 on fingertip 2 interacts bidentately with K77 and W78 on the ND helix. In contrast, in ActA, D406 points away due to a rotation of the ND helix, while in GDF8, these interactions are absent because a glycine occupies the equivalent position. In GDF11, the interactions remain preserved. This ionic lock binding strategy that engages conserved Asp on the fingertip 1 is likely important for outcompeting the interaction with positive residue in type I receptors at the K77 position (26, 27), as it is also employed by other antagonists (5, 10, 34, 35).

**Figure 3.**
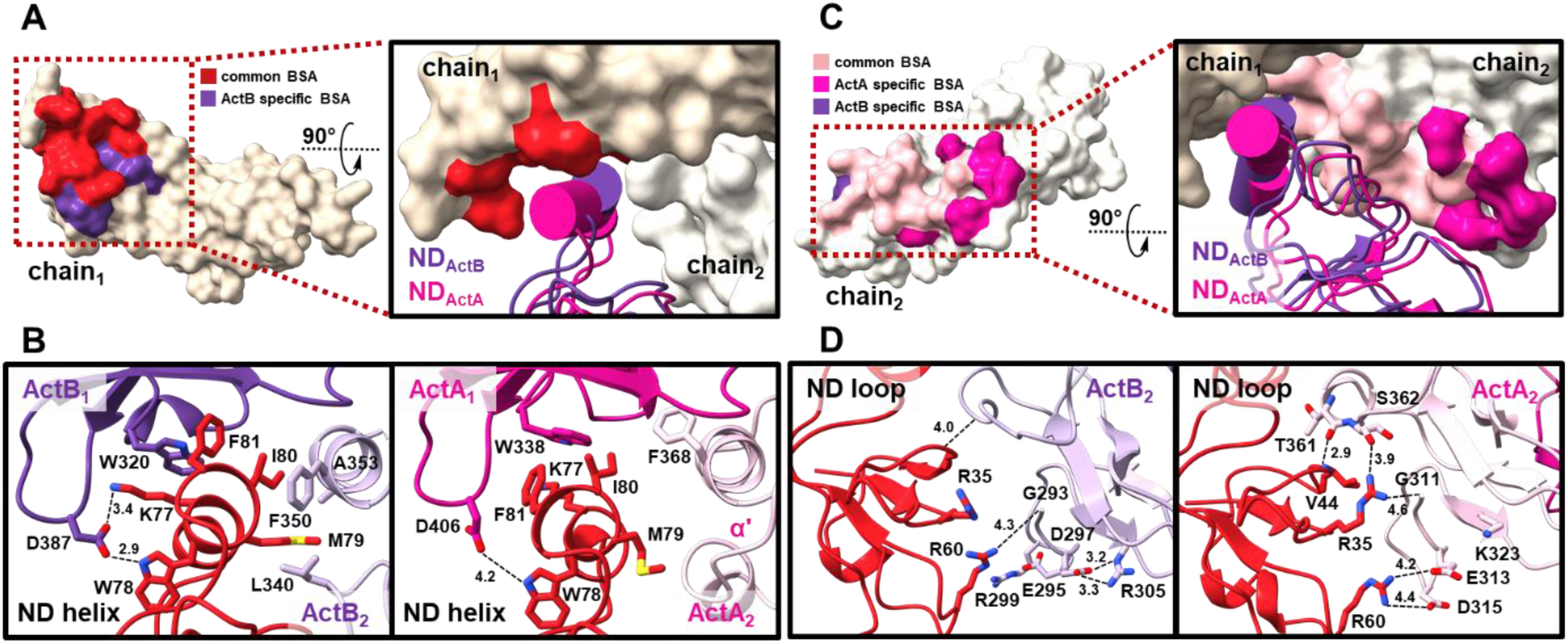
Type I Ligand Interface with the ND Domain. A) The buried surface area on chain_1_ by the ND that is shared between ActB and ActA is colored red, while the area specific to ActB is shown in purple. On the left, only the ActB chain_1_ is shown in beige surface representation. On the right, both chains and the ND helices are shown from a view rotated 90° horizontally. B) Detailed view of the ND helix binding site within the concave region of ActB (left) and ActA (right). C) The buried surface area on chain_2_ by the ND that is shared between ActB and ActA is colored pink, while the area specific to ActA is shown in magenta and to ActB in purple. On the left, only the ActA chain_2_ is shown in beige surface representation. On the right, both chains and the ND are shown from a view rotated 90° horizontally. D) Detailed view of the interaction between the ND loops and the N-terminus and wrist helix of ActB (left) and ActA (right).

On the other hand, ActB relies less on chain_2_ to interact with the ND than other activins (Fig. 3C). Specifically, arginines 35 and 60 on the ND loops, important for stabilizing the N-terminal region and α′ helix in ActA, do not engage in the same ionic interactions with ActB, but instead form intradomain hydrogen bonds (Fig. 3D). Compared to ActA and ActB, GDF8 and GDF11 possess two additional N-terminal residues that more efficiently stabilize ND loops via interactions with R35 and R60 (Fig. S7B). Furthermore, a hydrogen bond between ND V44/L45 and the ligand’s prehelix region, seen in ActA, GDF8, and GDF11 complexes, is missing in ActB, further supporting reduced engagement of ActB chain_2_ in ND binding. In computationally generated chimeras with swapped prehelix and wrist helix regions between ActA and ActB, we observed that in the ActB^A^:Fst288 complex, the α′ helix was formed as in wild type ActA, resulting in a 3.5 Å ND helix shift relative to the ActB:Fst288 crystal structure (Fig. S8). Interestingly, in the ActA^B^ mutant, where the prehelix region remained disordered, we did not notice the expected movement of the ND helix closer to the concave interface. Instead, only a slight shift at the base of the ND helix was detected, suggesting that, beyond steric hindrance, interactions with ligand fingers and fingertips also play an important role in ND positioning.

### Type II ligand epitope comprises a hydrophobic core and a polar periphery

The majority of the FSD1/FSD2 binding interface comprises numerous conserved and non-polar residues that form similar interactions across all four complexes and likewise engage the hydrophobic hotspot on type II receptors (23, 26, 27). E. g., a conserved leucine on the ligands’ finger 3 forms a CH···π interaction with Y188, which aligns with Y67 in BMPR2, F61 in ActRIIA, and Y60 in ActRIIB, while an isoleucine, a part of the conserved IAP motif in activins and BMPs, interacts with V190 (Fig. 4A). A valine at the same position is also preserved in Fstl3, which appears to be a mimicking strategy used by Fst-type antagonists to imitate V100 in ActRIIA and V99 in ActRIIB (23, 26). Unlike the central hydrophobic hotspot, surrounding peripheral residues show greater variability and are enriched in polar or charged amino acids. A key ionic interaction, also preserved in Fstl3 via R225 (5, 10), is formed between a conserved aspartate on fingertip 1 and R221 on FSD2. R221 is further stabilized by variable hydrogen bond-accepting residues on fingertip 2: Y389 in ActB, Q408 in ActA, E357 in GDF8, and Q389 in GDF11 (Fig. 4B). In the ActB complex, the conserved Asp also participates in a unique salt bridge with R194 on the FSD2 loop, which shifts the loop closer to the ligand compared to other activins (Δ_ActA_ = 5.5 Å, Δ_GDF8_ = 4.3 Å, Δ_GDF11_ = 3.4 Å). Unlike the ND helix interactions with conserved aspartates on the fingertip 1 that mimic type I receptors, the ionic contacts with FSD2 do not have direct counterparts in type II receptors, suggesting that these long-range interactions may have an important functional role in Fst binding.

**Figure 4.**
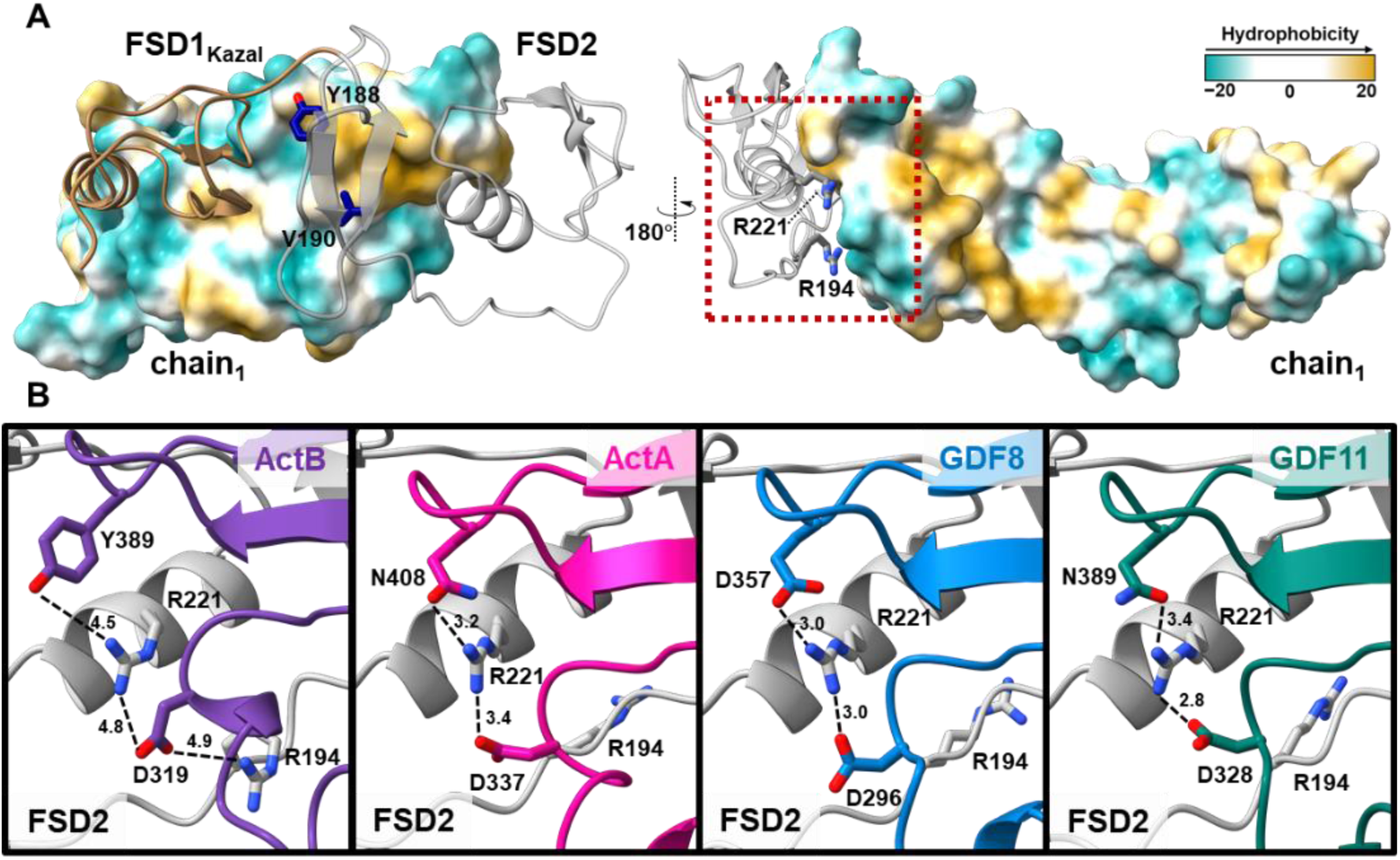
Type II Ligand Interface with the FSD1 and FSD2 Domains. A) Left: Surface representation of ActB chain_1_ colored according to hydrophobicity, showing hydrophobic contacts between the FSD1-FSD2 domains and the central patch on the knuckle epitope. Right: Front view of the FSD2 interaction with ActB fingers. The boxed region is shown enlarged in panel B. B) Detailed views of the interactions between R221 (and R194) in FSD2 and the ligand fingertips.

### FSD3 interacts with ActB fingertips at the expense of the other Fst288 molecule

Unlike in other complexes where FSD3 does not interact with ligand, in the ActB complex FSD3 engages with the uniquely acidic fingertip 2 of ActB (Fig. 5A). Specifically, R266, S268, and K298 on FSD3 form hydrogen or ionic bonds with E388. In ActA, the glutamate is replaced by a glycine, which cannot participate in similar interactions, resulting in a shift of FSD3 away from the ligand and toward adjacent Fst288. Due to its positioning closer to the ligand in the ActB complex, ND[1] lacks interactions with the FSD3[2] that are observed in the ActA complex, such as between R41 and E272 (Fig. 5B). In GDF8 and GDF11, the position equivalent to E388 in ActB is occupied by a lysine, whose interactions with hydrogen-donating residues on FSD3[2] might be even repulsive. Therefore, the Fst288 intermolecular interactions are not disrupted as in the ActB complex (Fig. S7C). Although analysis of evolutionarily related sequences shows that D387 and E388 in ActB fingertip 2 are variable (Fig. S9) (36), it appears that Fst288 has adapted its binding mode to accommodate this newly evolved feature in order to maximize the binding interaction.

**Figure 5.**
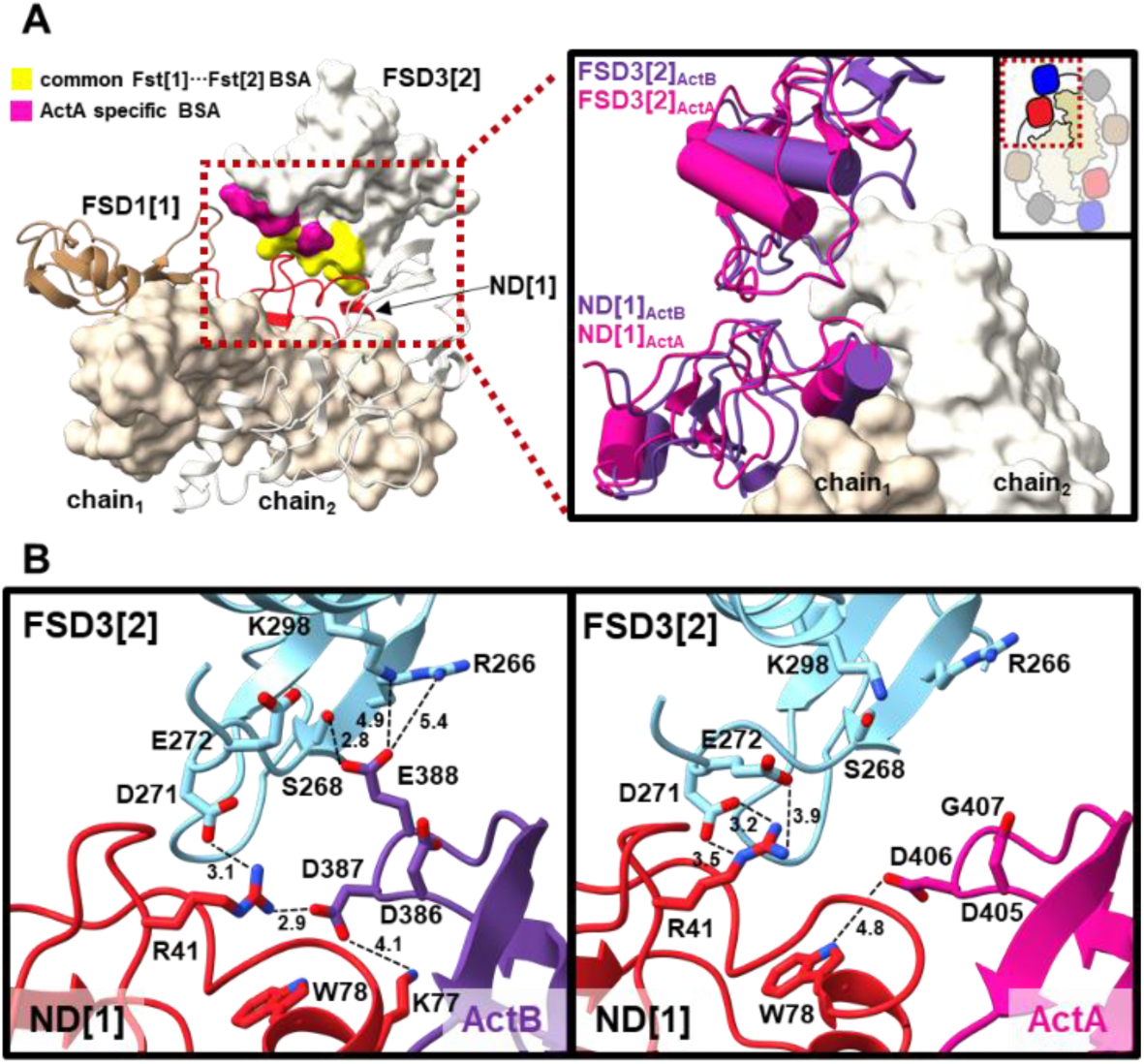
Interface between Ligands and Two Fst288 Molecules. A) On the left, the buried surface area on FSD3[2] by the Fst[1] that is shared between ActB and ActA is colored yellow, while the area specific to ActA is shown in magenta. ActA chain_1_ is shown in beige surface representation, chain_2_ in white cartoon, ND[1] in red, and FSD1[1] in brown. On the right, the ND and FSD3 domains from the ActB and ActA complexes are shown from a view indicated in the top right corner. B) Detailed view of the interactions between the FSD3[2] (light blue), ND[1] (red) and ligand in the ActB (left, in purple) and ActA (right, in pink) complexes.

### Computational Insight into Binding Energy Reveals Lack of Cooperativity in the ActB:Fst288 Complex

To assess the contribution of individual Fst288 domains to ligand binding, we employed MD simulations and the MM-GBSA model (Table S1). MM-GBSA enthalpy calculations revealed that two Fst288 molecules form the most stable complex with ActB, while the interaction enthalpies with ActA and GDF8 are more comparable (Fig. 6A). When total enthalpy is distributed across Fst288 domains, the ND binds more strongly to ActB than to ActA or GDF8 and is more important in the ActB binding than FSD2, while in the ActA and GDF8 complexes, ND and FSD2 contribute similarly (Fig. 6B). FSD1 consistently weakens binding across all three studied systems, as does FSD3 in the ActA and GDF8 complexes, but not in ActB. Furthermore, the interaction enthalpy between the two Fst288 molecules is highly favorable in the ActA and GDF8 complexes, but slightly unfavorable in the ActB (Fig. 6C). This suggests that interaction between two Fst288 is important for stabilizing the ActA and GDF8 complexes, whereas the ActB stability relies more on the individual affinities of each Fst288. This is consistent with previous studies that, based on the slope of luciferase reporter assay curves, suggested that cooperativity is prevalent in ActA and GDF8 antagonism (14). Our luciferase assays support this as well: while higher negative hill slope values in ActA and GDF8 indicate cooperativity, the value of −1.47 for ActB suggests a lack of cooperativity (Fig. 1C).

**Figure 6.**
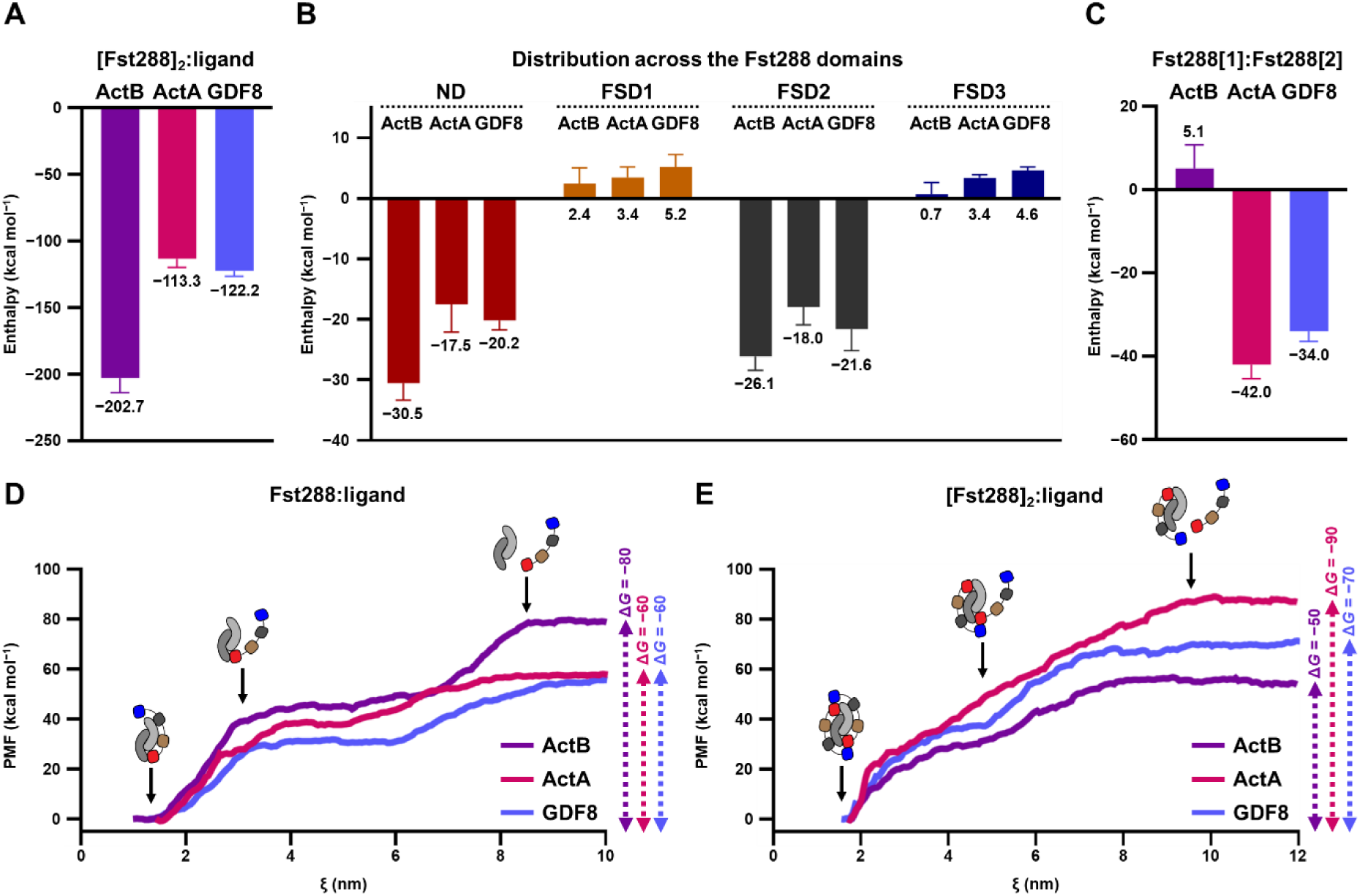
Computational Analysis of Interaction Enthalpy and Gibbs Free Energy in Activin:Fst288 Complexes. A) Calculated MM-GBSA enthalpy between two Fst288 molecules and ligands (in kcal mol^−1^). Values represent averages from five MD replicas. B) Distribution of calculated MM-GBSA enthalpy across individual Fst288 domains (in kcal mol^−1^). Values represent averages of both domains from five MD replicas. C) Calculated MM-GBSA enthalpy between the two Fst288 molecules (in kcal mol^−1^). Values represent averages from five MD replicas. Potential of mean force curves along the reaction coordinate, calculated from *umbrella sampling* simulations for dissociation of the first Fst288 from ligands D) or the second Fst288 from the Fst288:ligand complexes E), with corresponding Δ*G* values indicated on right (in kcal mol^−1^).

To further investigate cooperativity, we used an advanced sampling computational technique (*umbrella sampling*) which enables obtaining a free energy profile along a specific reaction coordinate, in this case dissociation of both Fst288 molecules. The calculated Gibbs free energy for the binding of the first Fst288 molecule is higher for ActB, than for ActA and GDF8 (Fig. 6D). All three energy profiles are characterized by two plateaus indicating biphasic binding: the initial rise reflects the dissociation of FSD1−3, while first plateau and the second increase correspond to the ND dissociation. While the first energy increase is similar for ActA and GDF8, and slightly higher for ActB, likely due to interactions with FSD3, the length of the first plateau and the final energy level are notably larger in ActB, suggesting that the ND binds most strongly to ActB. A similar pattern was observed in dissociation profiles of the ND alone (Fig. S10). The calculated Gibbs free energy for binding of the second Fst288 is highest for ActA, followed by GDF8, and lowest for ActB, confirming the importance of the intermolecular Fst288 interactions in the overall ActA and GDF8 affinity (Fig. 6E). With these additional interactions, the shape of curves changes, especially for ActA where the resulting monophasic profile indicates strong resistance against disruption of the Fst288···Fst288 interaction. The summed Δ*G* of both binding events align with previous reports that ActA (−150 kcal mol⁻¹) has higher affinity than ActB and GDF8 (−130 kcal mol⁻¹) (5, 6). However, similar effectiveness of Fst288 at inhibiting activins *in vitro* suggests that other interactors may be important in the cellular context.

### The ActB:Fst288 complex exhibits similar behavior to ActA:Fst288 in heparin affinity experiments

Fst288 is known to bind heparin and heparan sulfate (HS) via its heparin binding sequence (HBS) on FSD1 (Fig. 1A), which has implications in its localization on cell surface and endocytosis of ligands (37). The visualization of the electrostatic potential surface of complexes shows that GDF8 and GDF11 form a continuous electropositive crevice with Fst288 on the top, which likely binds heparin and HS (11, 12). In contrast, both ActA and ActB complexes lack such a localization of basic residues on the top surface (Fig. 7A) (8). To test how this affects the heparin binding, we used heparin affinity chromatography and computational approaches. In the chromatography assay, ActB:Fst288 eluted at a similar NaCl conductance as the ActA complex and Fst288 alone, whereas GDF11:Fst288 required a higher salt concentration to disrupt ionic interactions suggesting higher affinity (Fig. 7B,C).

**Figure 7.**
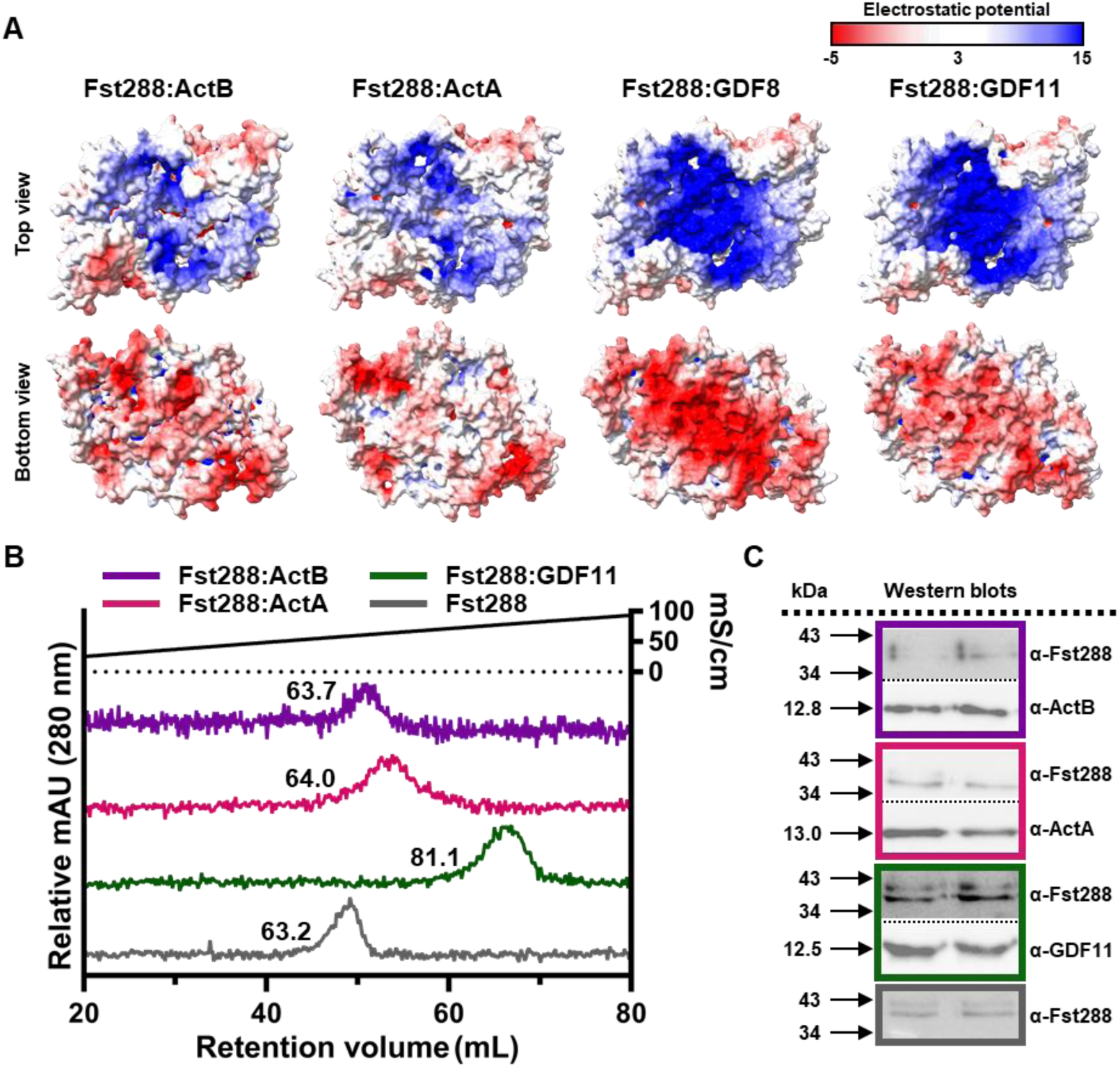
Interaction of Activin:Fst288 Complexes with Heparin. A) Top and bottom views of the electrostatic potential surfaces for Fst288:ligand complexes, calculated using APBS (Adaptive Poisson-Boltzmann Solver), with positive regions in blue and negative in red. B) UV absorbance traces from heparin affinity chromatography for Fst288 alone (gray), and for Fst288 complexes with ActB (purple), ActA (magenta), and GDF11 (green). Proteins were eluted using a NaCl gradient. Conductivity values are indicated at each peak maximum. C) Fractions from the heparin column for each sample (colored as in panel A) were analyzed by western blot using corresponding primary antibodies.

Using MD simulations and the MM-PBSA method, we calculated the interaction enthalpies between HS and Fst288 alone or in complexes. To examine the effect of chain length (degree of polymerization, dp), we simulated HS with dp8 (HS_8_) and dp16 (HS_16_), as differences in response began to appear for chain lengths greater than dp10 (37). Consistent with SPR data showing that longer heparin species form stronger interactions with Fst288 and its complexes (38), our calculations suggest higher affinity for HS_16_ than HS_8_, while when comparing complexes, GDF8 and GDF11 dominate over ActA and ActB (Fig. S11) (12). Furthermore, higher dp was associated with stronger stabilization of the Fst288 intermolecular interaction (Table S2), supporting that Fst288 may dimerize on the cell surface even before ligand binding (37). However, the Fst288 dimer with bound HS undergoes conformational changes to minimize the exposure of nonpolar surfaces to solvent, resulting in smaller width compared to ligand-bound complexes (Fig. S12). This likely influences only ligand association, as the interaction enthalpy between ligands and Fst288 was negligibly different than without HS (Table S2), consistent with kinetic studies showing that heparin/HS affected only the association rate, with no apparent effect on the dissociation rate (37).

Since ActB has an acidic fingertip 2 (D386, D387, E388) that interacts with FSD3, we speculated that mutating these residues to alanine could reduce ActB’s affinity for Fst288 and increase the complex’s affinity for HS. MD simulations showed that the triple mutant has a weakened interaction with Fst288 compared to WT, which allowed FSD3 to move closer to the HS, resulting in stronger stabilization (Table S2 and Fig. S13). This suggests that ActB fingertip 2, although important for Fst288 binding, may be responsible for reduced interaction with the extracellular matrix.

### Insight into the origin of Fst288 specificity

Beside activins, Fst288 binds moderately some BMP ligands, but does not antagonize the TGF-β subgroup (7). Sequence alignment of strongly antagonized activins with BMPs and non-antagonized TGF-β ligands shows that prehelix, wrist helix, and N-terminus − collectively forming the ND binding interface − differ substantially in both amino acid composition and length (Fig. S14). These regions are also the most variable between strongly antagonized activins and moderately antagonized BMPs. Since these differences do not correlate with Fst288’s ability to bind or antagonize ligands, we speculate that they fine-tune affinity rather than determine specificity. In contrast, the central part of the fingers in all TGF-β ligands, on both the concave and convex surfaces, contains conserved hydrophobic residues known to be important for receptors binding and likely responsible for stabilizing the complex upon Fst288 binding. This is supported by a computational analysis of individual residue contributions to the interaction enthalpy which shows the highest significance for these amino acids (Table S3 and S4). However, the fingertips and peripheral regions of both binding sites show considerable variability between ligands that bind Fst288 and those that do not. Namely, fingertip 1 in the binding ligands contains a conserved aspartate that interacts with R221 on FSD2, while the non-binding TGF-β subgroup has a positively charged lysine that likely acts repulsive. In binding ligands, this interaction is further supported by an acidic or hydrogen bond-accepting residue on fingertip 2, while TGF-βs possess a positive arginine or lysine at the same position (Fig. 4B and S14). Decomposition analysis shows that R221 is one of the major Fst288’s residues contributing to binding across all complexes (Table S3), and mutation of R221 in a truncated version of Fst288 abolishes binding to ActA (15), suggesting that R221 might play a key role in ligand recognition. Furthermore, in binding ligands, aspartates located on fingertip 2 interact with K77 and R41 in the ND, whereas in TGF-βs these interactions are prevented by the presence of glycine at the same position (Fig. S14). Interestingly, GDF8 also has G355, but the absence of a negative charge is compensated by a nearby glutamate (E357), which additionally stabilizes R221, as supported by decomposition analysis (Table S4). We therefore hypothesize that long-range ionic interactions between the exposed acidic fingertips of the ligands and positively charged residues in FSD2 and ND initiate binding and are responsible for Fst288 specificity.

## Discussion

Fst288 is a multidomain protein that functions as an endogenous antagonist of TGF-β ligands. In particular, it potently inhibits activins, only moderately some BMPs, but does not bind members of the TGF-β subgroup. Among activins, ActB is the least structurally characterized, and little is known about its regulation by Fst288, which limits our understanding of how Fst288 acts as a potent antagonist across the relatively diverse activin subgroup. To address this, we solved the crystal structure of the ActB:Fst288 complex and performed a comprehensive structural and computational comparison with other activin:Fst288 complexes.

From a global perspective, Fst288 binds activins in a similar manner, surrounding them with two molecules. By utilizing the flexibility of its interdomain loops, Fst288 accommodates ligands whose interchain angles can range from 50° (PDB: 1NYS) to 142° (PDB: 7U5O), and constrains them all to a similar conformation with an interchain angle of ∼110°. Strong binding is achieved by mimicking structural features of receptors: e.g. the ND helix positions itself in the ligand’s concave surface where the universal type I receptor motif would normally bind, while the EGF domain of FSD2 imitates the three-finger toxin fold of type II receptors on the knuckle epitope. In addition to these secondary-structure similarities, Fst288 also forms receptor-like interactions with the ligands, ensuring high affinity and effectively outcompeting receptor binding.

Despite the overall similarity, a deeper structural analysis reveals certain peculiarities in the ActB binding to Fst288 compared to other activins. At the type I receptor interface, which in the Fst288 complex is occupied by the ND domain, the fingers of chain_1_ and the N-terminus and wrist helix of chain_2_ in ActB are equally engaged, whereas in other complexes ligand chain_2_ dominates. While the type II receptor interface appears similar across all complexes, another striking difference occurs at the FSD3 interface involving the ligand and the ND of the adjacent Fst288 molecule. Namely, unlike in other complexes where FSD3[1]···ND[2] contacts between two Fst288 are more extensive, in the ActB complex FSD3 instead interacts with the uniquely acidic ligand fingertip 2. Calculations of the interaction enthalpy suggest that these differences result in more extensive contacts between Fst288 and ActB, giving the highest value relative to ActA and GDF8. Further computational analysis revealed that ND binds ActB most strongly, which negatively impacts its ability to engage in interactions with the second Fst288 and indirectly contribute to the overall affinity through cooperative effects. FSD3 interactions with ActB additionally limit the potential for cooperativity, as evidenced by the unfavorable interaction enthalpy between the two Fst288 in the ActB complex.

This is further supported by *umbrella sampling* calculations of the free energy profiles for the dissociation of the first and second Fst288 molecules from their ligand complexes. These curves allow the calculation of Gibbs free energy which, by including the entropy term, more accurately correlate with experimentally measured *K*_d_ values: Fst288 has a 10-fold higher affinity for ActA than for ActB and GDF8 (5, 6). Binding of the first Fst288 to ActB is most favorable due to the highest interaction enthalpy. However, binding of the second molecule is more favorable in the ActA and GDF8 complexes than in ActB, and in fact becomes more favorable than the first binding event, indicating a cooperative effect. In contrast, binding of the second Fst288 to ActB is less favorable than the first, likely due to enthalpy–entropy compensation. The enthalpic gain from strong binding can be offset by a loss in entropy, as the interacting partners become more conformationally restricted. In other words, the tighter and more specific the interaction, the less entropically favorable it becomes. This effect is most evident in ActB, whose apo state is substantially more flexible than that of ActA or GDF8, as shown by MD simulations. Thus, while the binding of the first Fst288 to ActB is primarily enthalpically driven, still allowing a certain degree of freedom, the second binding event incurs a larger entropic penalty as the ligand becomes locked into a more rigid conformation. The summed Δ*G* values reflect this trend, with ActA exhibiting the highest total affinity (−150 kcal mol⁻¹), while ActB and GDF8 show comparable but lower values (−130 kcal mol⁻¹). The importance of cooperativity for ActA binding was shown in experiments when Fstl3 ND was replaced with that of Fst288 which reduced affinity for ActA; however, attaching FSD3 to that construct restored the binding, indicating that ND···FSD3 contacts enhance the ND ability to effectively bind ActA (10). Therefore, it is likely that Fst288’s broad inhibitory activity among activins arises from its ability to adjust its binding mode and utilize cooperativity to enhance complex stability when individual domains are insufficient for high-affinity binding. From a functional perspective, achieving highly stable complexes and complete ligand neutralization may serve to prevent the formation of subfunctional 1:1 antagonist:ligand assemblies that could act as receptor sinks and exert broader effects beyond the activin subgroup.

Despite their different affinities, an *in vitro* luciferase reporter assay yielded similar inhibition curves and nearly identical nanomolar IC₅₀ values for all activins. This suggests that, in a cellular context factors beyond simple binding affinity may influence the antagonist’s overall effectiveness. Fst288 is known to bind components of the extracellular matrix and localize to the cell surface, which may facilitate the endocytosis of Fst-bound ligands (39, 40). Previous studies have shown that, depending on the electrostatic potential on the ligand’s surface, Fst288 complexes may have altered affinities for heparin and heparan sulfate (HS). E.g. GDF8 and GDF11, which have a localized patch of positively charged residues on their top surface, form Fst288 complexes with higher heparin affinity than Fst288 alone (11, 12). In contrast, in our heparin affinity experiments and computational analyses, ActB behaved similarly to ActA, whose Fst288 complexes bind heparin comparable to Fst288 alone, consistent with the absence of distinct positively charged areas on the ligands. Another factor that may affect binding to HS is the ligand:Fst288 interaction itself. E.g. the acidic fingertip 2 of ActB which interacts with FSD3 may prevent it from extending toward HS, which is typically anchored in the top crevice between the two HBS regions. Computationally introducing a triple alanine substitution at residues 387–389 slightly destabilized the ligand–Fst288 complex but increased HS affinity, suggesting the importance of fingertip 2 for the regulation of ligand by the antagonist. Together, these data suggest that ligand-specific features and the interaction strength, as well as the distribution of electrostatic potential and binding to the extracellular matrix, contribute to the overall effectiveness of antagonists.

Finally, to understand why Fst288 binds activins and BMPs but not TGF-βs, we compared their sequences. The type II receptor-binding site features hydrophobic residues mostly conserved in both binding and non-binding ligands, whereas the type I epitope includes residues that vary widely among binding ligands, suggesting that these amino acids alone cannot determine Fst288 specificity. The most striking difference appears at the fingertips where all the binding ligands contain acidic residues, while TGF-βs carry positively charged amino acids, implicating their role in specificity. Based on these observations, together with our computational and previous experimental findings showing that heparin and HS promote Fst288 dimerization in the absence of a ligand (37), we propose a mechanism in which a cell surface-localized Fst288 dimer recognizes ligands through long-range ionic interactions involving charged residues on their exposed edges (Fig. 8). As MD simulations suggest, Fst288 dimerization is driven by ionic contacts between FSD3 and the HBS on FSD1 which must be disrupted by even stronger ionic interactions with ligands for binding to occur. Following initial electrostatic recognition, short-range hydrophobic contacts at the type II epitope confer high affinity, while engagement of the highly variable prehelix and wrist helix regions by the ND domain primarily functions to fine-tune affinity and ultimately determine the stability of the antagonist–ligand complexes.

**Figure 8.**
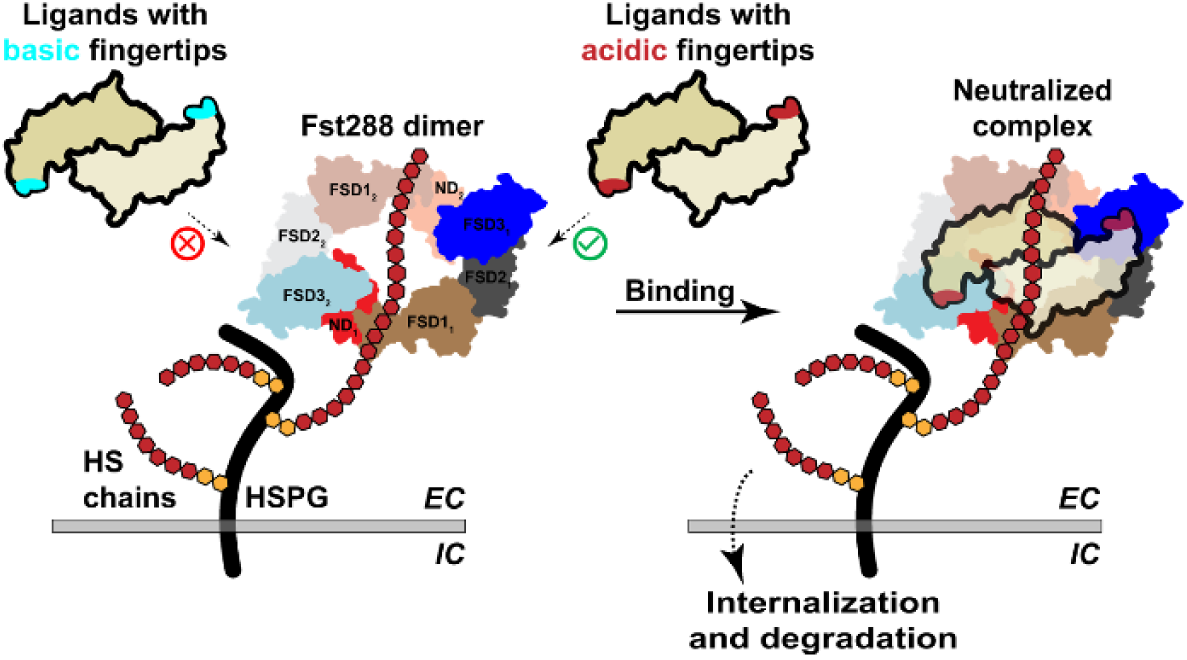
Proposed Mechanism for Neutralization of Ligands by Fst288. The first step involves recognition through ionic interactions, followed by the formation of short-range hydrophobic interactions that are responsible for high affinity and complex stability. Abbreviations: HSPG = Heparan Sulfate Proteoglycan, EC = extracellular, IC = intracellular.

Our study, through a comprehensive comparison of activin:Fst288 complexes, provides insights into the broad activity of Fst288 and its specificity for the activin subgroup. While the details of ligand binding to Fst288 offer a structural basis for the future design of inhibitors with desired selectivity to target conditions with elevated ligand levels, a deep understanding of Fst288 function may also help elucidate how other endogenous antagonists regulate TGF-β signaling.

## Data availability

The activin B:Fst288 structure has been deposited in PDB under ID: 9Z3T. Software information: AlphaFold3 (https://alphafoldserver.com/), AMBER22 and AmberTools23 (https://ambermd.org/), CCP4 suite (https://www.ccp4.ac.uk/), ConSurf (https://pubmed.ncbi.nlm.nih.gov/36718848/), Coot (https://www2.mrc-lmb.cam.ac.uk/personal/pemsley/coot/), GROMACS 2022.4 (https://www.gromacs.org/), HDOCK (http://hdock.phys.hust.edu.cn/), MODELLER (https://salilab.org/modeller/), Phenix (https://www.phenix-online.org/), PROPKA3.1(https://server.poissonboltzmann.org/pdb2pqr), UCSF Chimera 1.17.3(https://www.cgl.ucsf.edu/chimera/). Data to reproduce the theoretical calculations are available from LH or TBT upon request.

## Supporting information

This article contains supporting information.

## Acknowledgments

Funding for the work was provided by a grant to TBT NIH R35 GM134923. LH thanks the Advanced Research Computing (ARC) center at the University of Cincinnati, Cincinnati, OH, for providing computing resources.

## Author contributions

LH, RGW, EJG, and TBT designed research; LH, RGW, JAH, JFM, CK, and EJG performed research; LH, RGW, JAH, and EJG analyzed data; and LH wrote the paper.

## Abbreviations

ActA: Activin A
ActB: activin B
BMP: bone morphogenetic protein
Fst288: follistatin 288
FSD1–3: follistatin domains 1–3
HS: heparan sulfate
HBS: heparin binding sequence
MDS: molecular dynamics simulations
MM-GBSA: Molecular Mechanics-Generalized Born Surface Area
MM-PBSA: Molecular Mechanics-Poisson-Boltzmann Surface Area
ND: N-terminal domain
PMF: potential of mean force
PCA: principal component analysis
RMSD: root mean square deviation
SPR: surface plasmon resonance
TGF-β: transforming growth factor-β

## Supporting Information

## Experimental and computational procedures

### Protein purification and complex generation

Individual proteins were produced and purified as previously described (8, 11, 12). All the experiments were performed with proteins produced and purified by the authors. Complexes were formed by adding ligands to an excess of Fst288 at a 1:2.5 molar ratio, incubated at 4°C for few hours and purified on a Superdex 200 column (Amersham Biosciences).

### ActB:Fst288 complex crystal structure determination

Purified ActB:Fst288 complex was concentrated to 6.6 mg mL^−1^ in 0.02 M TRIS pH 7.5 and 0.05 M NaCl before it was mixed 1:1 in a hanging drop experiment with a solution containing 0.15 M ammonium sulfate, 0.1 M HEPES and 15% (w/v) PEG 3350. Diffraction experiments were performed at the Argonne National Laboratory Advanced Photon Source 23ID beamline and processed as previously described (11, 12). Phasing was performed by molecular replacement using Phaser (41) and the CCP4 suite (42), using an AlphaFold3 (43) model of a complex containing a human ActB dimer and two Fst288 molecules. Iterative refinement was carried out with Phenix (44) and Coot (45). Coordinates have been deposited in the PDB (PDB ID: 9Z3T).

### Luciferase reporter assay

The luciferase reporter assay using HEK293-(CAGA)_12_ cells (originally derived from RRID:CVCL_0045) was performed as previously published (5, 12, 14). Cells were seeded in 96-well plates and cultured for 24 hours. The growth medium was then removed and replaced with serum-free medium + 0.1% BSA containing ligands at a concentration of 0.62 nM, which was mixed with twofold serial dilutions of Fst288. After 18 hours, the cells were lysed, and luminescence was measured using a Synergy H1 Hybrid plate reader (BioTek). Activity data were imported into GraphPad Prism 8 and analyzed using nonlinear regression with a variable slope to calculate the IC₅₀.

### Heparin affinity analysis

Heparin affinity was determined as previously published (11, 12). Namely, 100 μg of Fst288, either alone or in complex with ActB, ActA, or GDF11, was applied to a 1 mL HiTrap column (Amersham Biosciences) and eluted with a linear gradient of up to 2 M NaCl over 120 column volumes.

### Computational analysis

#### 1. Molecular dynamics (MD) simulations

The resolved structure of the ActB:Fst288 complex, along with previously determined structures of Fst288 complexes with ActA (8), GDF8 (11), and GDF11 (12) were used as starting models for MD simulations. Because of the high similarity between GDF8 and GDF11, most analyses were performed on the GDF8 MD simulations; however, MD simulations of the GDF11 complex with heparan sulfate were also examined to compare the results with the heparin affinity chromatography data. Missing residues were modeled using Chimera’s Modeller plugin (46), and the protonation states of ionizable amino acid residues were estimated by PROPKA 3.1 (47). Complexes were solvated in 10 Å-octahedral boxes, neutralized with corresponding number of counterions, and parametrized using the AMBER ff19SB force field for proteins and the OPC model for water (48, 49). Systems were subjected to geometry optimization in AMBER22 with periodic boundary conditions applied in all directions (50). The optimized systems were gradually heated from 100 to 298 K over 1 ns under constant volume and then equilibrated for 7 ns at a constant pressure, followed by productive, unconstrained MD simulations lasting 300 ns. Starting positions of ligand:Fst288:HS complexes were generated using HDOCK server (51), while ActB^A^ (featuring swapped prehelix and wrist helix regions: 335-362 residues in ActB were replaced by 353-380 residues in ActA) and ActA^B^ (featuring swapped prehelix and wrist helix regions: 353-380 residues in ActA were replaced by residues 335-362 in ActB) chimeras were modeled in complex with Fst288 using AlphaFold3 (43). Apo ligand structures were obtained by removing Fst288 from the complexes, whereas the Fst288 dimer was generated by deleting GDF8 from the corresponding complex structure. The same MD simulation protocol described above was applied to all systems. Trajectory processing and data analysis were performed using the CPPTRAJ module implemented in AMBER (52). For enthalpy calculations of ligand:Fst288 and ligand:Fst288:HS complexes, five or three replicas of 50 ns MD simulations were run, respectively. Interaction enthalpies of ligand:Fst288 complexes were computed from 3,000 frames taken from the final 40 ns of each simulation using the MM-GBSA protocol and decomposed into *per-residue* contributions (53). To better account for the highly negative charge of HS in the ligand:Fst288:HS complexes, binding enthalpies were calculated using the MM-PBSA method with an internal dielectric constant set to 6, as previously reported (54).

#### 2. Umbrella sampling simulations

Umbrella sampling simulations of ActB:Fst288, ActA:Fst288, and GDF8:Fst288 complexes were performed using GROMACS 2022.4 (55), employing periodic boundary conditions in all directions. The systems were set up in 10.0 nm x 14.0 nm x 25.0 nm rectangular boxes, solvated using the TIP3P water model, and neutralized with the appropriate number of counterions. Each system was oriented so that Fst288 was aligned with the z-axis, along which it was pulled out by a harmonic spring with force constant of 3,000 kJ mol^−1^ nm^−2^, moving away from the center of mass of the ligand or ligand:Fst288 complex at constant velocity of 0.01 nm ps^−1^. To prevent the rest of the system from being displaced along with Fst288, a harmonic position restraint with a force constant 5,000 kJ mol^−1^ nm^−2^ was imposed on the wrist helices of the ligands. For the subsequent windows simulations, snapshots were extracted from the pulling simulations such that the z-component of the ligand···Fst288 or ligand:Fst288···Fst288 distance increased by 0.2 nm between neighboring windows, accounting for approximately 50 windows in total. For each window, a 20 ns MD simulation was performed using a harmonic force constant of 2,000 kJ mol^−1^ nm^−2^. If necessary, additional windows using a force constant of 3,000 kJ mol^−1^ nm^−2^ were added to ensure sufficient overlap between histograms. Potential of mean force (PMF) curves were calculated using the Weighted Histogram Analysis Method (WHAM) with the *gmx wham* tool in GROMACS (56).

---

**Figure S1.**
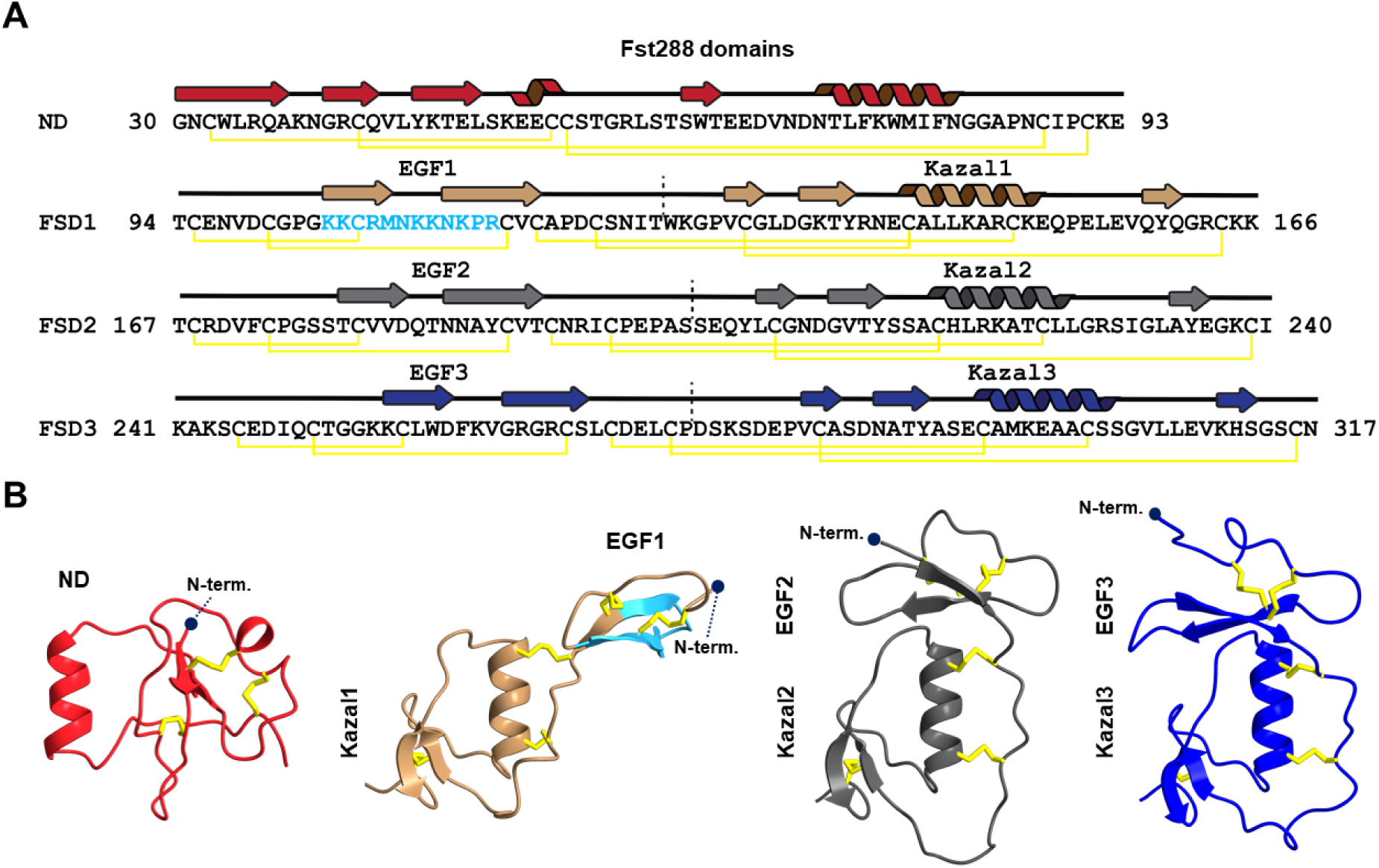
Domain Organization of Fst288. A) Sequence of four Fst288 domains with the secondary structure shown above. Heparin binding sequence (HBS) in FSD1 is given in cyan. B) Structures of individual Fst288 domains from the ActB:Fst288 complex (PDB ID: 9Z3T) aligned through the Kazal1 helix, with connecting disulfide bonds indicated.

**Figure S2.**
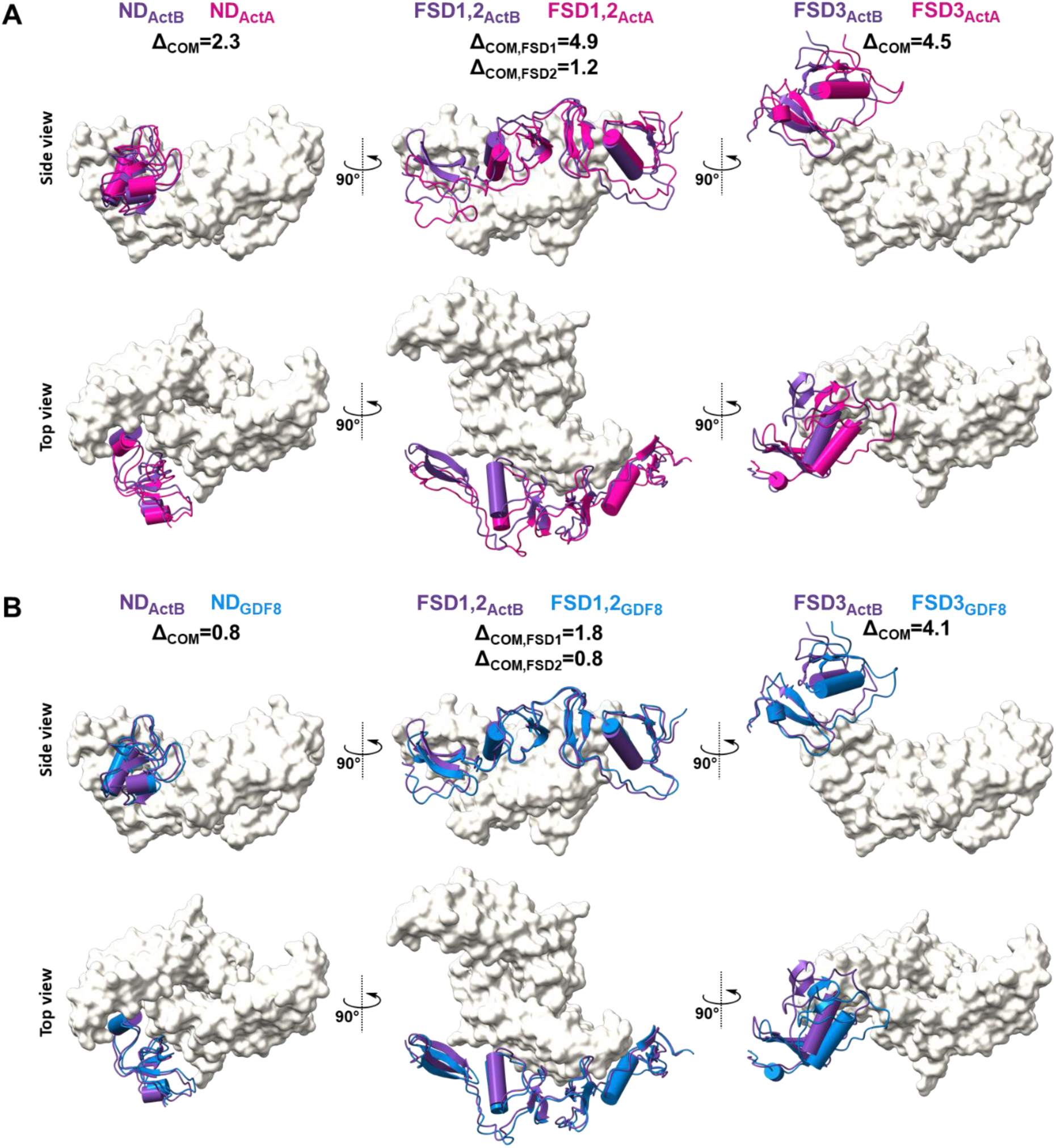
Structural Comparison of Individual Fst288 Domains in Complexes with Activins. Position of the Fst domains in the A) ActA and B) GDF8 complexes relative to ActB aligned through ligand chain_1_. Fst domains are colored purple in the ActB complex, magenta in ActA, and blue in GDF8. Differences in the centers of mass of individual domains relative to those in the ActB complex are given in Å. Only the ActB ligand is shown, in the beige surface representation.

**Figure S3.**
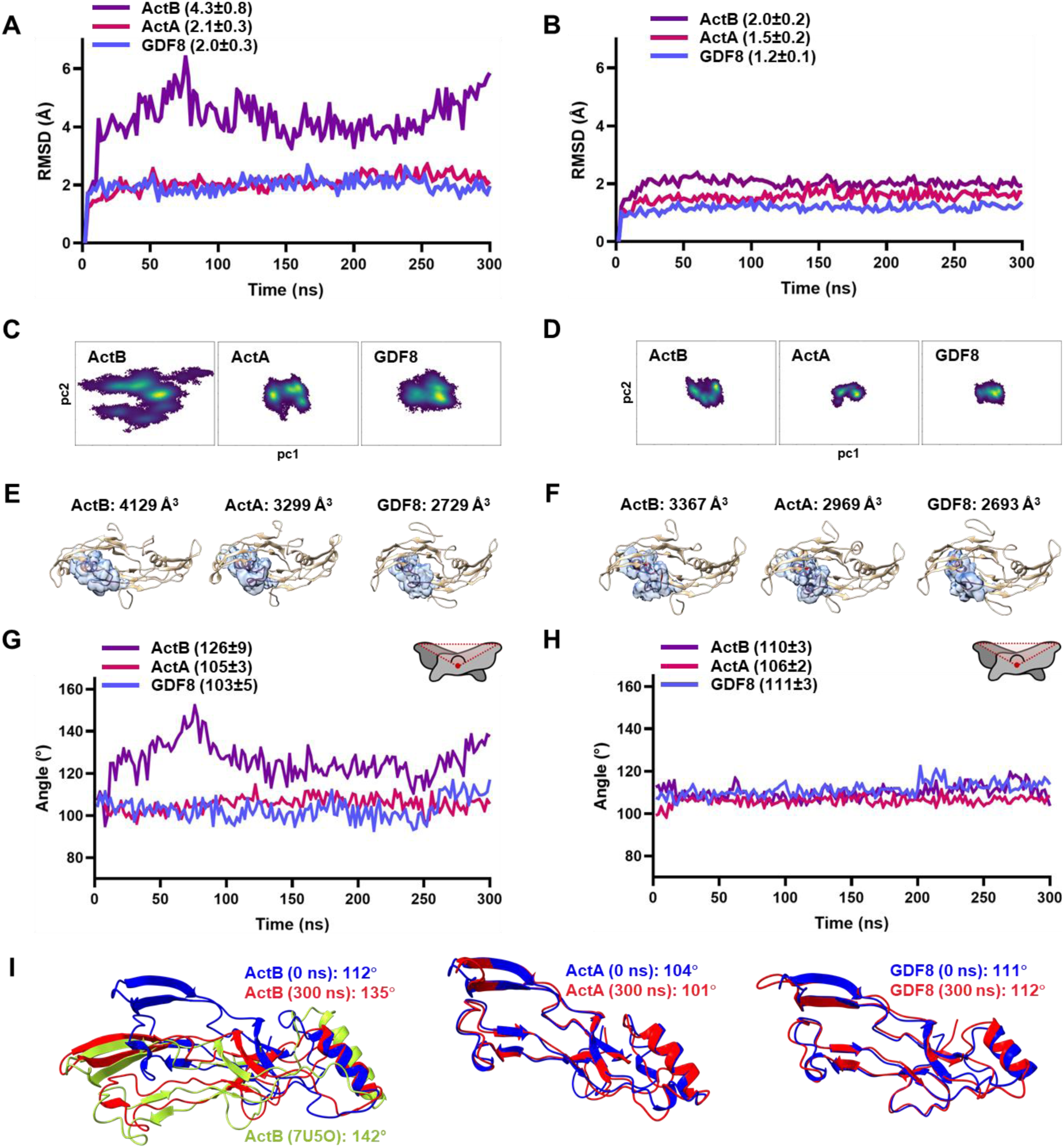
Analysis of Ligand Flexibility in Apo and Complex MD Simulations. RMSD graphs for backbone atoms of ligands in MDS of their A) apo forms and B) complexes with Fst288. Principal Component Analysis (PCA) results for ligands from MDS of their C) apo forms and D) complexes with Fst228, shown as density plots of the first two principal components. The density of points is indicated by a color gradient: yellow represents higher density, and dark blue represents lower density. 3D histograms showing the positions of the prehelix and wrist helix regions of ligands during MDS of their E) apo forms and F) complexes with Fst228, together with the average structures for reference. For clarity, the occupied volume of only one chain is shown (in blue), however, the calculated values represent averages from both chains. Interchain angle in MDS of G) apo forms and H) complexes measured between V392_1_-C372_1_-V392_2_ in ActB, I410_1_-C390_1_-I410_2_ in ActA, and I359_1_-C339_1_-I359_2_ in GDF8. I) Superposition of the starting and final frames of the apo MDS for ActB (left), ActA (middle), and GDF8 (right). On the left, ActB from the ActRIIB complex (PDB: 7U5O) is shown in light green. Ligands are aligned through one chain (not shown) to illustrate changes in the interchain angle.

**Figure S4.**
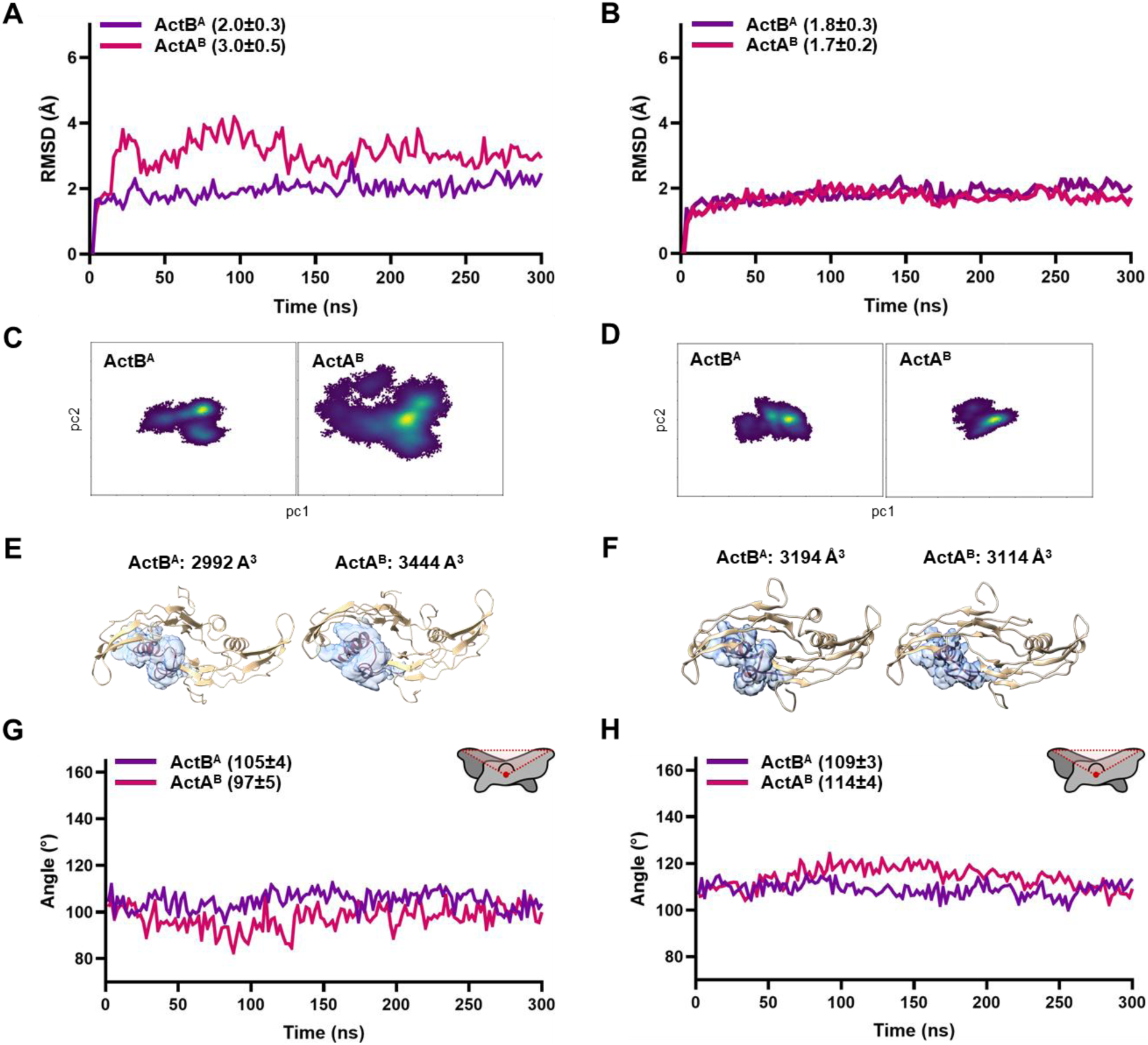
Analysis of Chimeric Ligands Flexibility in Apo and Complex MD Simulations. RMSD graphs for backbone atoms of ActB^A^ and ActA^B^ mutants in MDS of their A) apo forms and B) complexes with Fst288. Principal Component Analysis (PCA) results for ActB^A^ and ActA^B^ mutants from MDS of their C) apo forms and D) complexes with Fst228, shown as density plots of the first two principal components. The density of points is indicated by a color gradient: yellow represents higher density, and dark blue represents lower density. 3D histograms showing the positions of the prehelix and wrist helix regions of ActB^A^ and ActA^B^ mutants during MDS of their E) apo forms and F) complexes with Fst228, together with the average structures for reference. For clarity, the occupied volume of only one chain is shown (in blue), however, the calculated values represent averages from both chains. Interchain angle in MDS of G) apo forms and H) complexes measured between V392_1_-C372_1_-V392_2_ in ActB^A^, and I410_1_-C390_1_-I410_2_ in ActA^B^.

**Figure S5.**
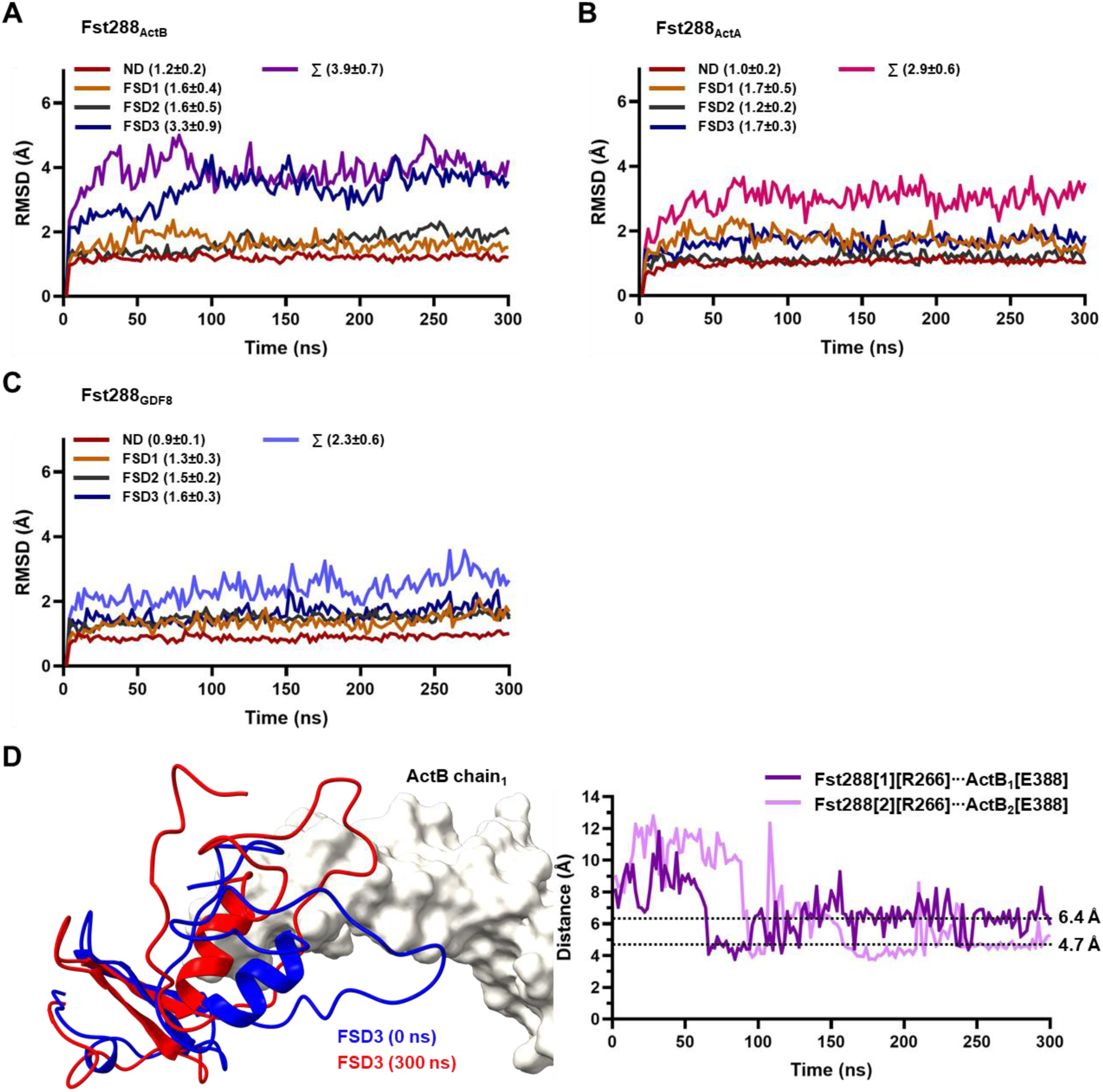
Analysis of Fst288 Flexibility in MD Simulations of the Activin Complexes. RMSD graphs for backbone atoms of Fst288 domains in complexes with A) ActB, B) ActA, and C) GDF8. D) Left: Position of FSD3 relative to ActB at the beginning (blue) and end (red) in MD simulation of the AcB:Fst288 complex. Right: Distance graphs between Fst288[R266-CZ] and ActB[E388-CD] on both chains over the course of the simulation.

**Figure S6.**
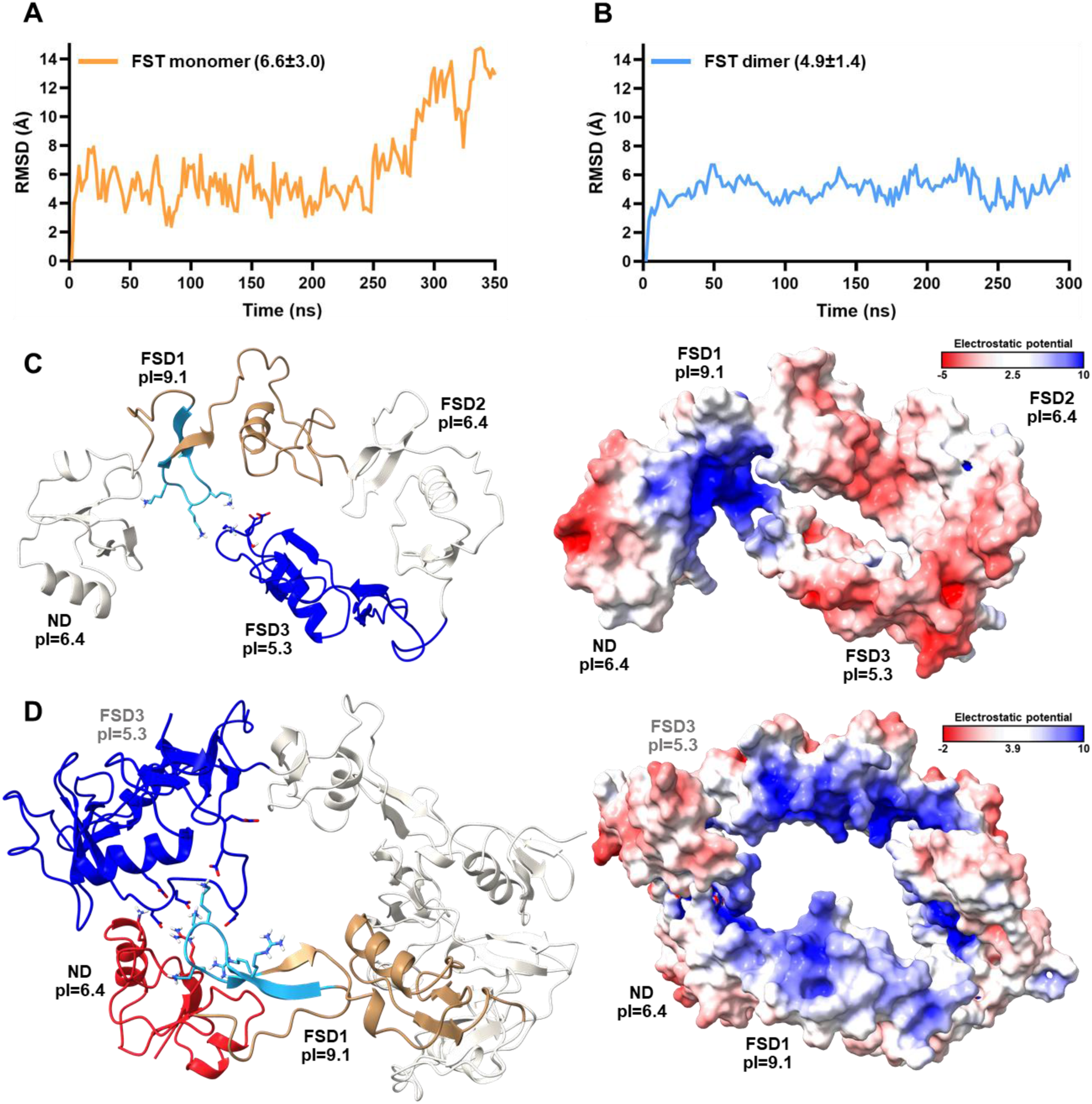
Analysis of Fst288 Flexibility in Apo MD Simulations. RMSD graphs for backbone atoms of Fst288 in A) single-molecule MD simulation and B) MD simulation of a complex of two Fst288 molecules, both in the absence of ligands. C) On the left: final frame of the apo MD simulation of a single Fst288 molecule showing the interaction between positively charged residues in the heparin binding sequence (HBS) of FSD1 and negatively charged residues in FSD3. FSD1 is colored light brown with the HBS in light blue, and FSD3 is shown in dark blue. On the right: electrostatic potential surface of a single Fst288 molecule after 300 ns calculated using APBS (Adaptive Poisson-Boltzmann Solver), with positive regions in blue and negative regions in red. D) On the left: final frame of the apo MD simulation of two Fst288 molecules showing the interaction between positively charged residues in the HBS of FSD1 from one molecule and negatively charged residues in FSD3 of the other. The ND is colored red, FSD1 is light brown with HBS in light blue, and FSD3 is dark blue. On the right: electrostatic potential surface of two Fst288 molecules after 350 ns calculated using APBS, with positive regions in blue and negative regions in red.

**Figure S7.**
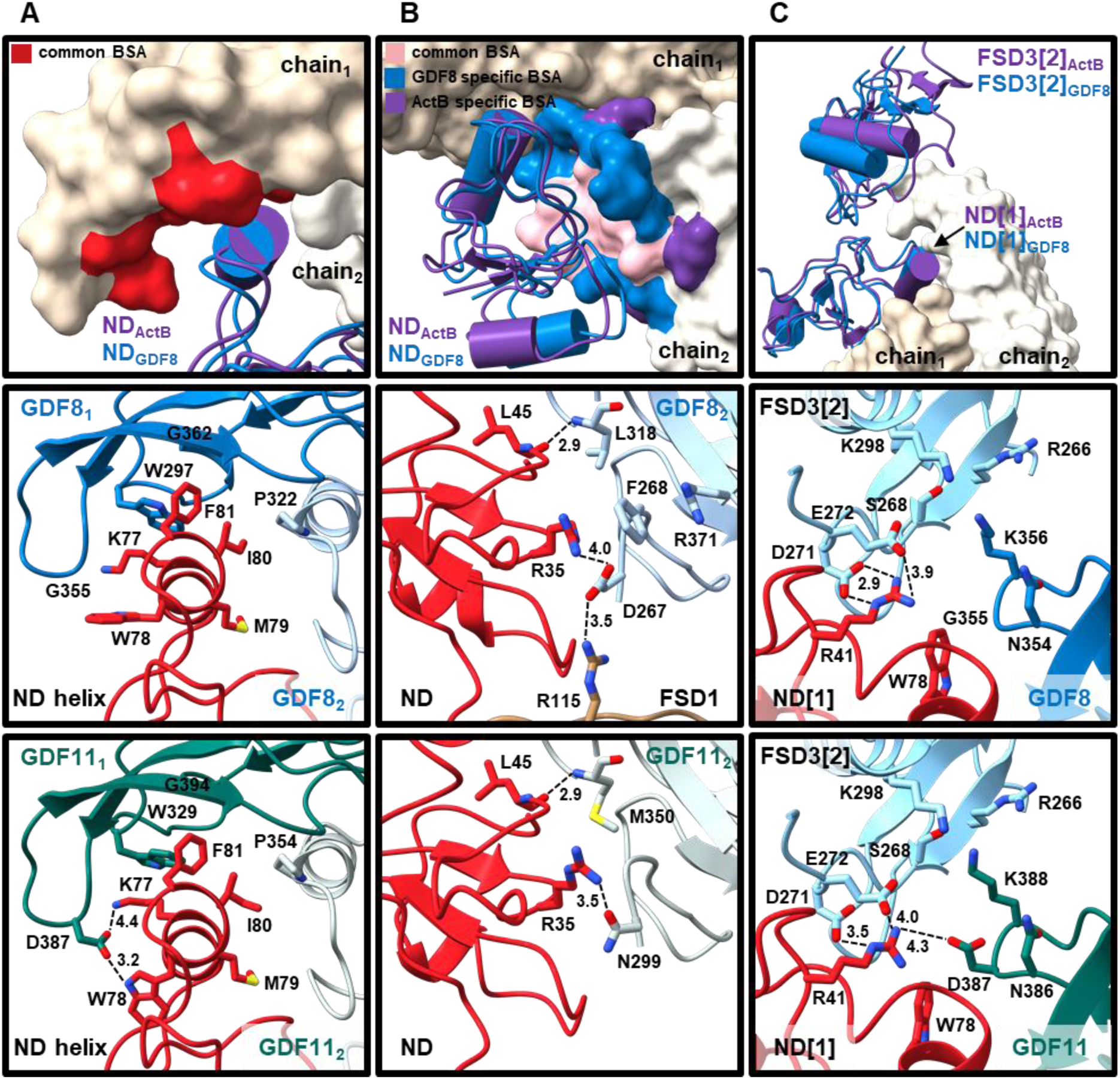
Type I and Fst[1]···Fst[2] Interfaces in GDF8 and GDF11 Complexes. A) Top: The buried surface area on chain_1_ by the ND that is shared between ActB and GDF8 is colored red. ActB chain_1_ is shown in beige and chain_2_ in white surface representation. Middle and bottom: Detailed view of the ND helix binding site within the concave region of GDF8 and GDF11. B) Top: The buried surface area on chain_2_ by the ND that is shared between ActB and GDF8 is colored pink, while the area specific to GDF8 is shown in blue and to ActB in purple. Middle and bottom: Detailed view of the interaction between the ND loops and the N-terminus and wrist helix of GDF8 and GDF11. C) Top: The ND and FSD3 domains from the ActB and GDF8 complexes. Middle and bottom: Detailed view of the interactions between FSD3[2], ND[1] and the ligand in the GDF8 and GDF11 complexes.

**Figure S8.**
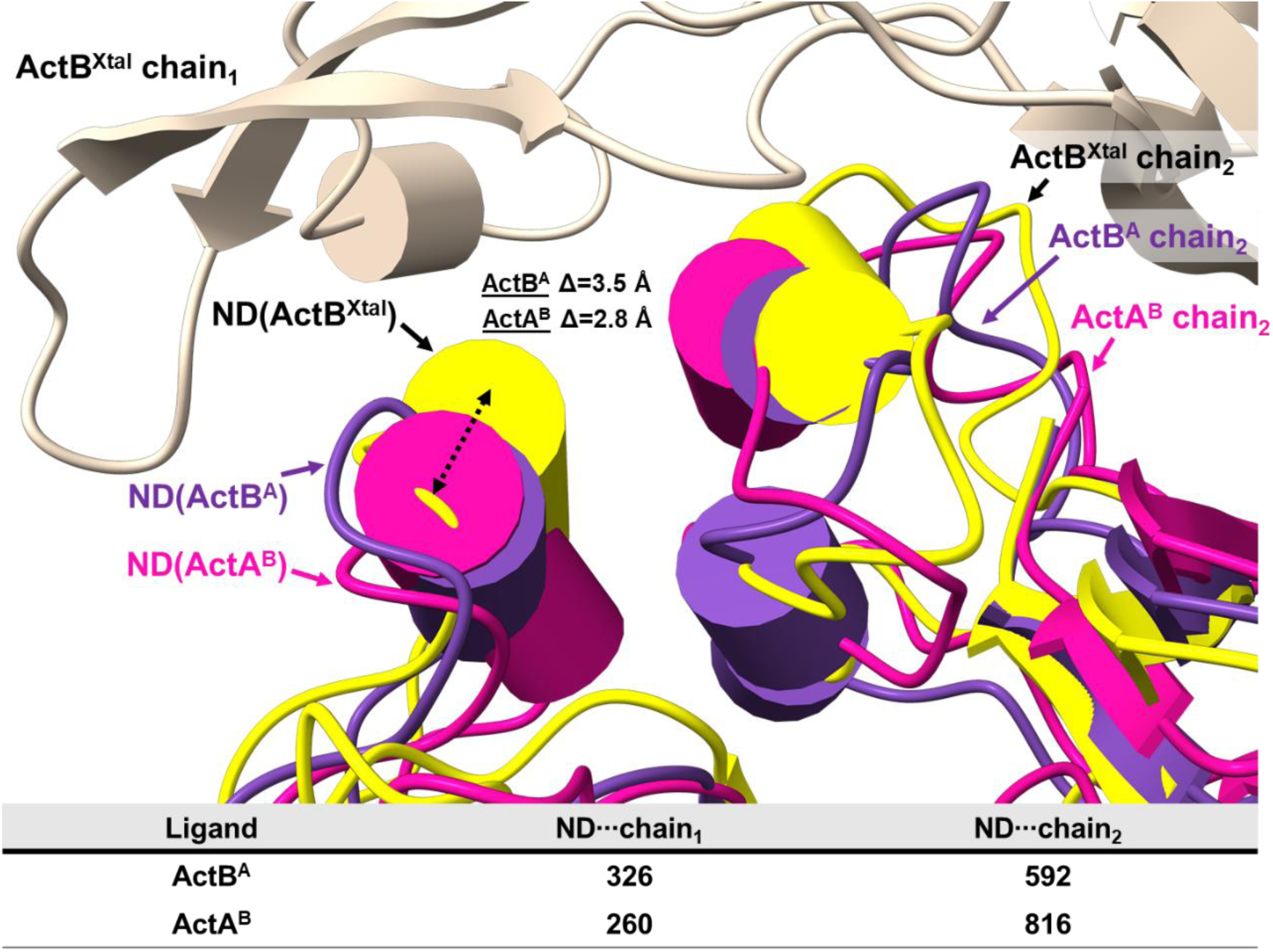
Type I Interface of Chimeric Ligands with the ND Domain. Superposition of the final frames from MD simulations of Fst288 complexes with ActB and ActA mutants (with swapped prehelix and wrist helix regions) onto the crystal structure of ActB:Fst288. Alignment was performed using chain_1_, and only chain_1_ from the crystal structure is shown. The ND helix and chain_2_ from the ActB crystal structure are colored yellow, from the ActB^A^ mutant in purple, and from the ActA^B^ mutant in magenta. Calculated BSA values (≥15 Å²) for ND interactions with ActB^A^ and ActA^B^ chains are given in the table below.

**Figure S9.**
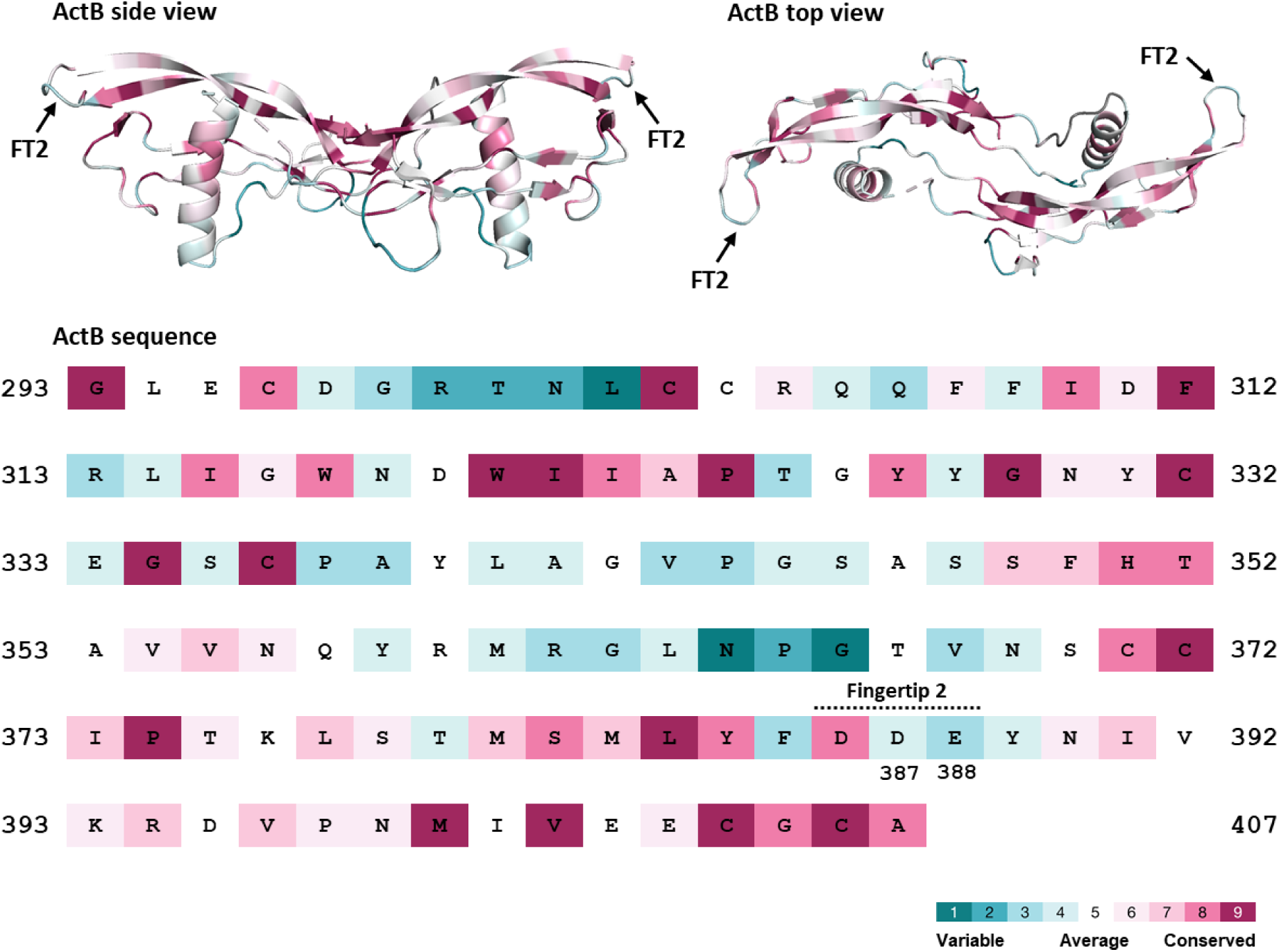
ConSurf Evolutionary Conservation Profile of ActB. The ActB structure is shown in side and top views (top), and the corresponding sequence is displayed below with conservation scores for each residue, colored according to the indicated scheme.

**Figure S10.**
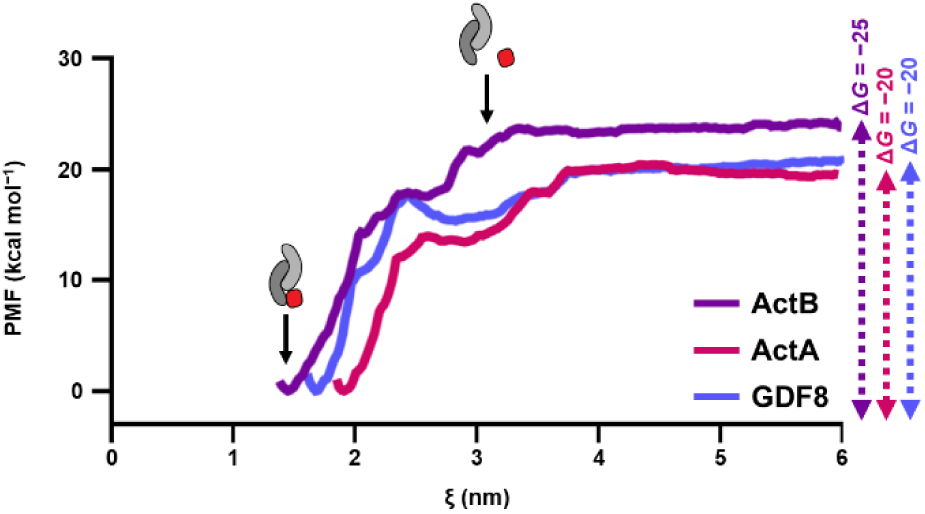
Computational Evaluation of Gibbs Free Energy between Activins and the ND Domain. Potential of mean force curves along the reaction coordinate, calculated from *umbrella sampling* simulations for dissociation of the ND from ligands, with corresponding Δ*G* values indicated on right (in kcal mol^−1^).

**Figure S11.**
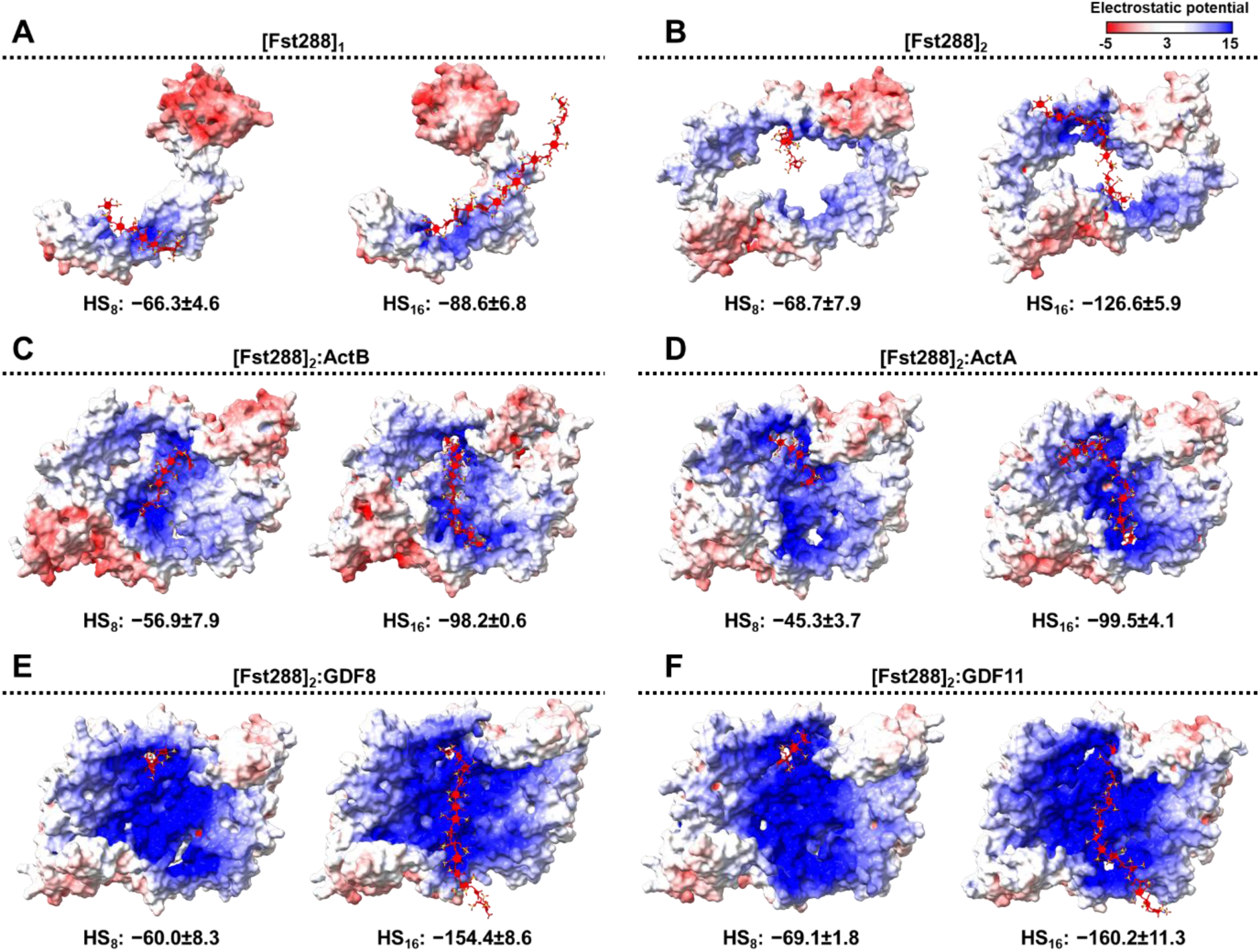
Interaction Enthalpy of Fst288 and Activin:Fst288 Complexes with Heparan Sulfate. Top views of electrostatic potential surfaces calculated using APBS (Adaptive Poisson-Boltzmann Solver) for: A) monomeric Fst288, B) dimeric Fst288, and C–F) Fst288 complexes with ActB, ActA, GDF8, and GDF11, respectively, each with bound heparan sulfate of 8 or 16 degrees of polymerization. Positive electrostatic regions are shown in blue and negative regions in red. Interaction enthalpies between Fst288 alone or Fst288:ligand complexes and heparan sulfate (HS) were calculated using the MM-PBSA method and are given in kcal mol⁻¹.

**Figure S12.**
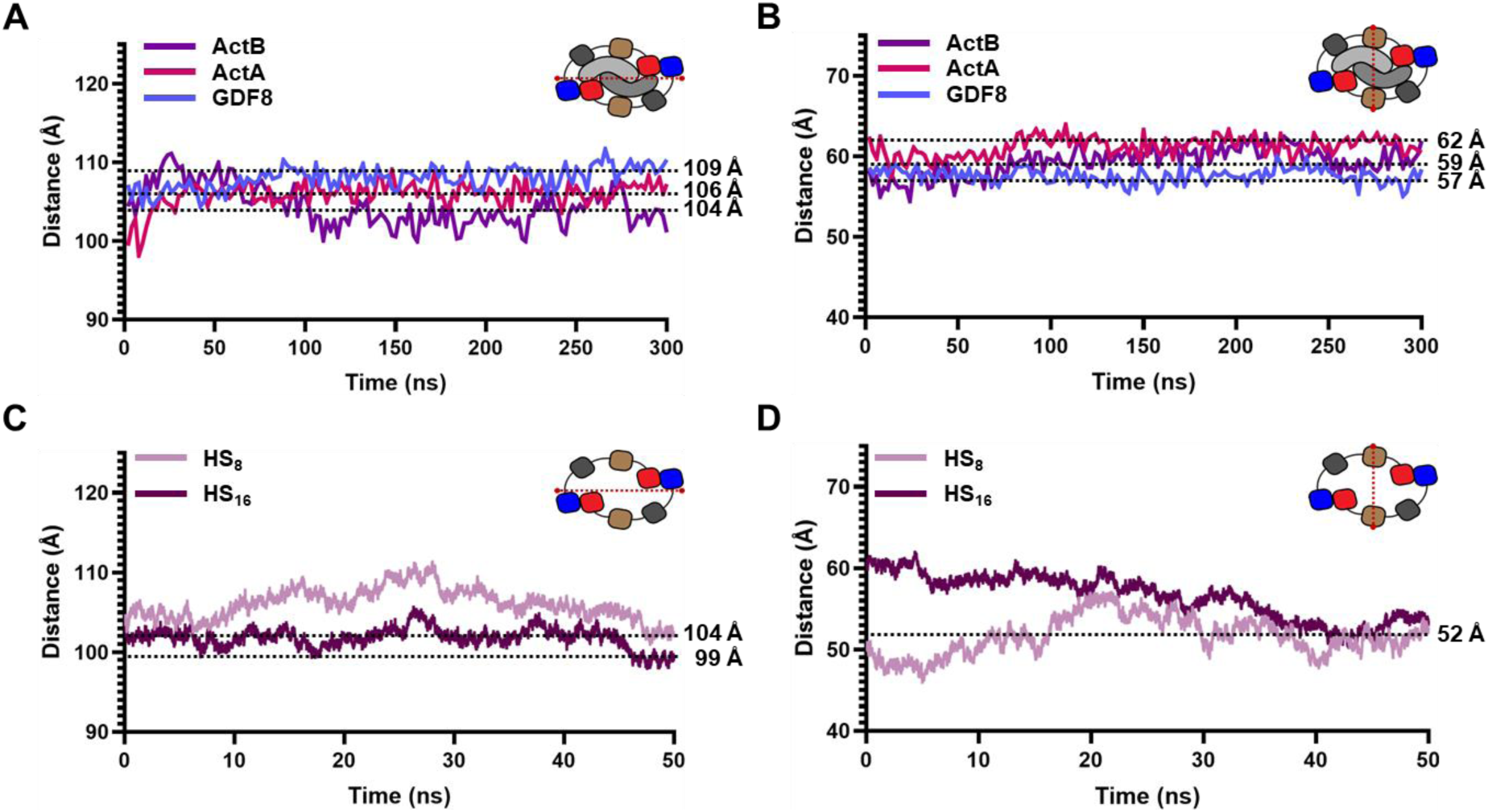
Comparison of Fst288 Dimer Dimensions in Complex with Activins or Heparan Sulfate. A) Overall complex length measured between K243_1_-K243_2_ in MD simulations of ActB, ActA, and GDF8 complexes. B) Overall complex width measured between C106_1_-C106_2_ in MD simulations of ActB, ActA, and GDF8 complexes. C) Length of the complex of two Fst288 measured between K243_1_-K243_2_ in MD simulations with HS_8_ and HS_16_. D) Width of the complex of two Fst288 measured between C106_1_-C106_2_ in MD simulations with HS_8_ and HS_16_.

**Figure S13.**
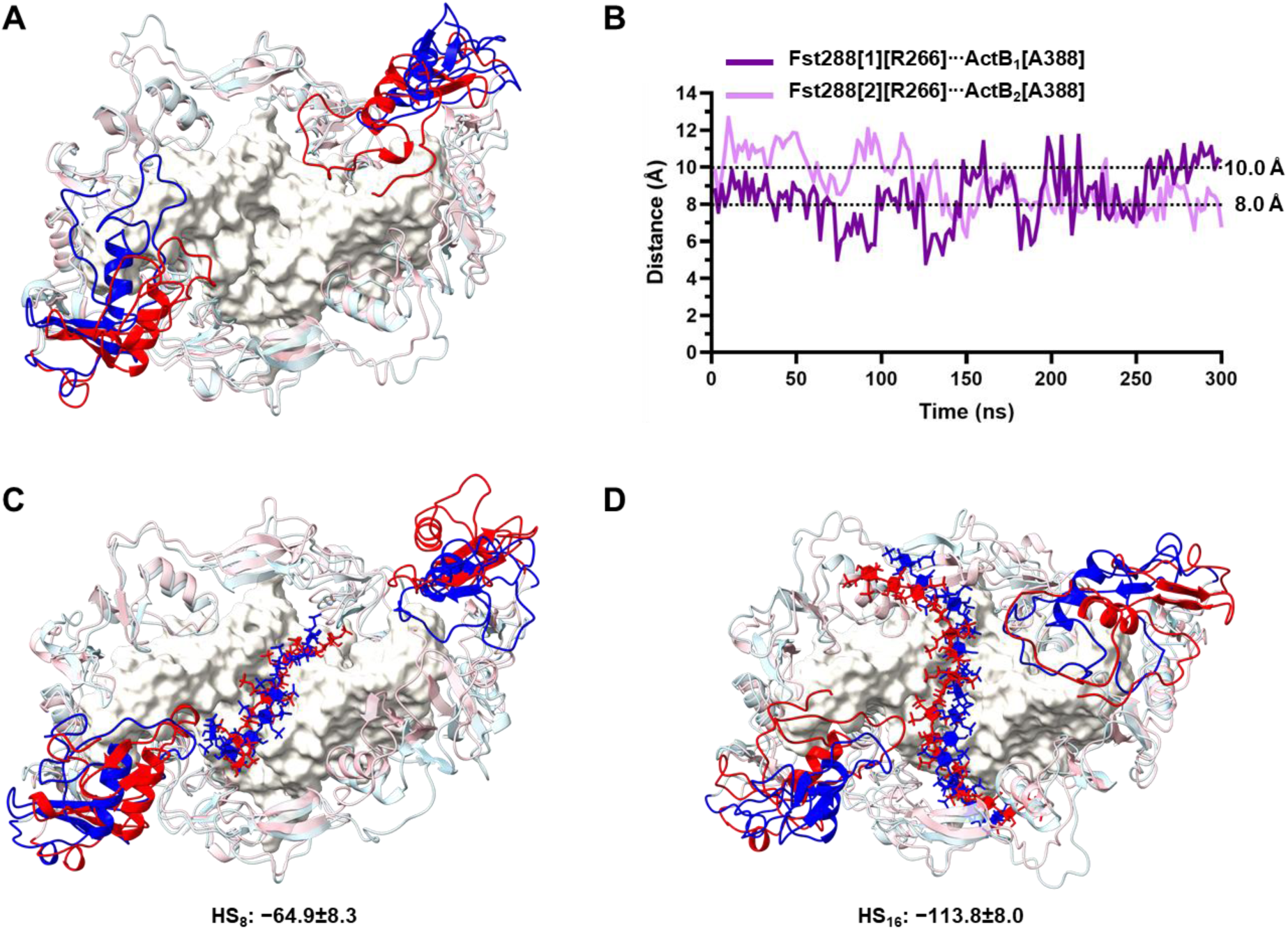
Interaction of the ActB D^386^A/D^387^A/E^388^A Triple Mutant with Fst288 or with Fst288 and Heparan Sulfate. Top views of the Fst288 complex with the ActB D^386^A/D^387^A/E^388^A triple mutant: A) without heparan sulfate (HS), C) with HS_8_, and D) with HS_16_. Each mutant complex is superimposed with the corresponding wild-type Fst288:ActB complex. In the mutant complex, FSD3 and HS are colored red, and the remaining Fst288 domains are pink. In the wild-type complex, FSD3 and HS are colored blue, and the remaining Fst288 domains are light blue. B) Distance plots between Fst288[R266-CZ] and the triple mutant ActB[A388-CB] on both ligand chains over the course of the simulation without HS.

**Figure S14.**
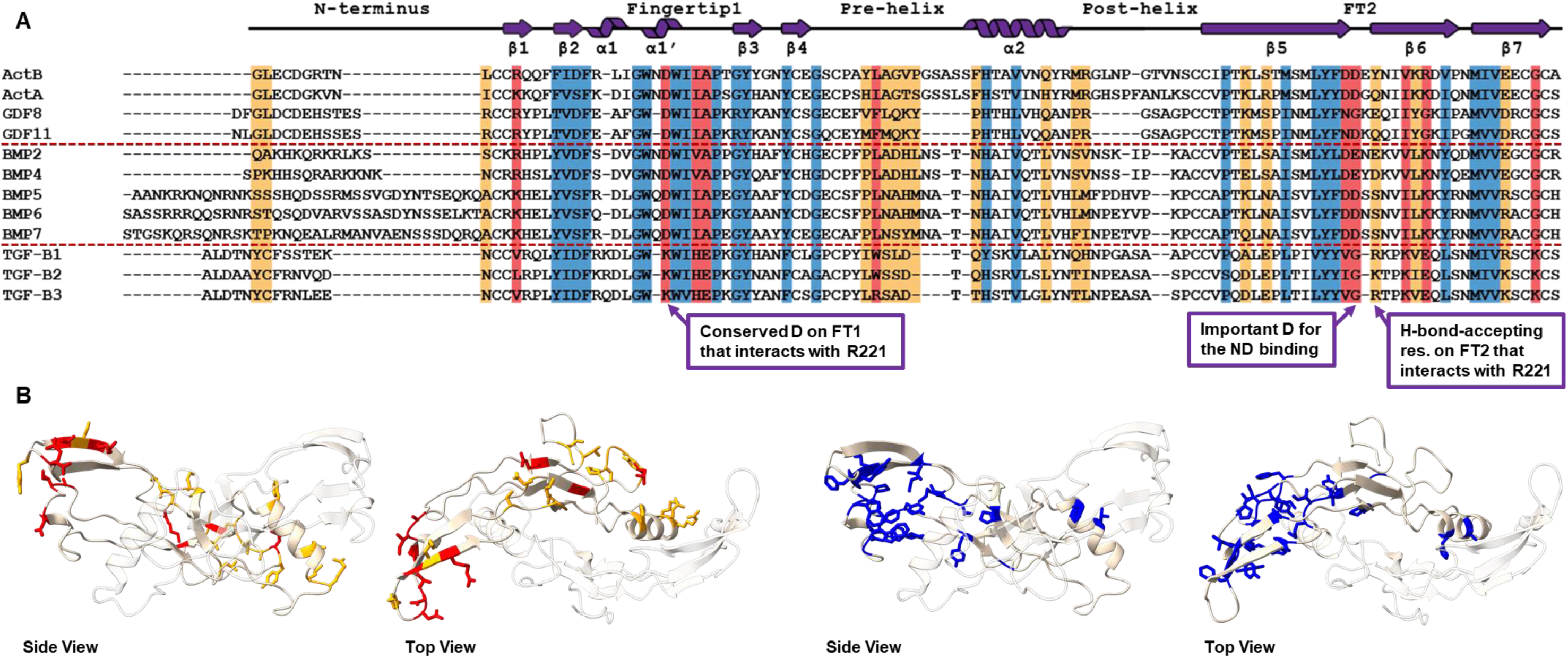
Insight into the Origin of Fst288 Specificity through Sequence Alignment of TGF-β Ligands. A) Sequence alignment of the strongly antagonized activins, moderately antagonized BMPs, and non-binding TGF-β ligands. Conserved amino acids among all ligands are highlighted in blue, residues that differ between antagonized and non-binding ligands are shown in red, and those that differ between the strongly and moderately antagonized ligands are shown in orange. The secondary structure of ActB is shown at the top of the alignment. B) On the left, the red and orange residues, and on the right, the blue residues from the sequence alignment are mapped onto the ActB structure in both side (left) and top (right) views.

**Table S1.** A) Calculated MM-GBSA enthalpy between the two Fst288 molecules and ligands with values from five MD replicas. B) Distribution of calculated MM-GBSA enthalpy across individual Fst288 domains with values of both domains from five MD replicas. C) Calculated MM-GBSA enthalpy between the two Fst288 molecules with values from five MD replicas.

| | $\Delta H\{[Fst288]_2:ligand\}$ (kcal mol<sup>-1</sup>) | | | | | | |
| --- | --- | --- | --- | --- | --- | --- | --- |
|  | #1 | #2 | #3 | #4 | #5 | AVG | STD |
| **ActB** | -191.7 | -206.4 | -202.3 | -219.3 | -194.1 | **-202.7** | 9.8 |
| **ActA** | -108.7 | -112.9 | -119.2 | -120.0 | -105.6 | **-113.3** | 5.7 |
| **GDF8** | -118.6 | -128.9 | -121.9 | -122.3 | -119.2 | **-122.2** | 3.7 |

B)
| Distribution of $\Delta H\{[Fst288]_2:ActB\}$ across Fst288 domains (kcal mol<sup>-1</sup>) | | | | | | | |
| --- | --- | --- | --- | --- | --- | --- | --- |
|  | #1 | #2 | #3 | #4 | #5 | AVG | STD |
| ND | -28.4 | -26.87 | -33.90 | -31.85 | -28.42 | **-30.5** | 2.6 |
|  | -31.25 | -32.03 | -26.71 | -34.48 | -31.51 |  |  |
| FSD1 | 0.48 | 0.07 | 5.28 | -0.45 | -0.69 | **2.4** | 2.5 |
|  | 4.60 | 2.03 | 5.67 | 1.73 | 5.64 |  |  |
| FSD2 | -25.95 | -29.03 | -28.50 | -25.74 | -21.87 | **-26.1** | 2.2 |
|  | -25.03 | -23.9 | -28.58 | -27.61 | -25.26 |  |  |
| FSD3 | -0.81 | 0.16 | 1.80 | 0.57 | 1.73 | **0.7** | 1.8 |
|  | 1.64 | 1.45 | -4.10 | 2.36 | 1.89 |  |  |
| Distribution of $\Delta H\{[Fst288]_2:ActA\}$ across Fst288 domains (kcal mol<sup>-1</sup>) | | | | | | | |
|  | #1 | #2 | #3 | #4 | #5 | AVG | STD |
| ND | -12.92 | -16.32 | -18.69 | -10.04 | -12.18 | **-17.5** | 4.4 |
|  | -21.38 | -17.47 | -22.80 | -20.01 | -23.19 |  |  |
| FSD1 | 3.50 | 3.44 | 3.66 | 4.72 | 3.92 | **3.4** | 1.7 |
|  | 4.21 | 4.04 | -1.53 | 3.89 | 4.21 |  |  |
| FSD2 | -19.09 | -21.64 | -16.72 | -18.19 | -18.92 | **-18.0** | 2.8 |
|  | -19.91 | -13.39 | -12.67 | -19.13 | -20.28 |  |  |
| FSD3 | 3.47 | 4.23 | 3.41 | 3.95 | 3.86 | **3.4** | 0.5 |
|  | 2.55 | 2.98 | 2.76 | 3.25 | 3.14 |  |  |
| Distribution of $\Delta H\{[Fst288]_2:GDF8\}$ across Fst288 domains (kcal mol<sup>-1</sup>) | | | | | | | |
|  | #1 | #2 | #3 | #4 | #5 | AVG | STD |
| ND | -20.97 | -21.66 | -19.27 | -20.34 | -17.22 | **-20.2** | 1.5 |
|  | -22.30 | -21.24 | -19.46 | -18.49 | -20.86 |  |  |
| FSD1 | 3.73 | 5.72 | 8.76 | 6.14 | 7.02 | **5.2** | 2.0 |
|  | 4.17 | 2.78 | 6.50 | 2.06 | 4.64 |  |  |
| FSD2 | -22.85 | -20.34 | -28.25 | -14.21 | -21.00 | **-21.6** | 3.4 |
|  | -19.54 | -23.11 | -20.32 | -23.71 | -22.48 |  |  |
| FSD3 | 4.96 | 4.31 | 4.14 | 3.87 | 4.90 | **4.6** | 0.5 |
|  | 4.97 | 5.51 | 4.25 | 4.23 | 5.20 |  |  |

C)
| | $\Delta H\{Fst288[1]:Fst288[2]\}$ (kcal mol<sup>-1</sup>) | | | | | | |
| --- | --- | --- | --- | --- | --- | --- | --- |
|  | #1 | #2 | #3 | #4 | #5 | AVG | STD |
| **ActB** | 10.1 | 12.3 | -0.5 | 2.5 | 1.2 | **5.1** | 5.1 |
| **ActA** | -41.4 | -40.6 | -37.4 | -46.0 | -44.5 | **-42.0** | 3.0 |
| **GDF8** | -38.0 | -34.4 | -32.5 | -33.3 | -31.6 | **-34.0** | 2.2 |

**Table S2.** Calculated MM-PBSA interaction enthalpies between the two Fst288 molecules and ligands in the absence of heparan sulfate (HS), and in the presence of HS with degrees of polymerization 8, and 16.

| | $\Delta H[\text{Fst288:Fst288}] \text{ (kcal mol}^{-1}\text{)}$ | | |
| --- | --- | --- | --- |
|  | Without HS | HS<sub>8</sub> | HS<sub>16</sub> |
|  | -9.0±0.4 | -11.6±2.8 | -17.5±3.4 |

| | $\Delta H\{[\text{Fst288}]_2:\text{ligand}\} \text{ (kcal mol}^{-1}\text{)}$ | | |
| --- | --- | --- | --- |
|  | Without HS | HS<sub>8</sub> | HS<sub>16</sub> |
| **ActB** | -103.0±2.9 | -106.0±7.4 | -106.6±5.1 |
| **ActB<sup>387-389→Ala</sup>** | -85.0±5.6 | -90.5±4.3 | -79.5±6.1 |
| **ActA** | -81.5±8.0 | -85.4±7.0 | -87.6±3.2 |
| **GDF8** | -83.0±8.2 | -82.7±4.7 | -96.9±7.1 |
| **GDF11** | -68.0±3.1 | -70.2±2.3 | -70.5±5.6 |

**Table S3.**
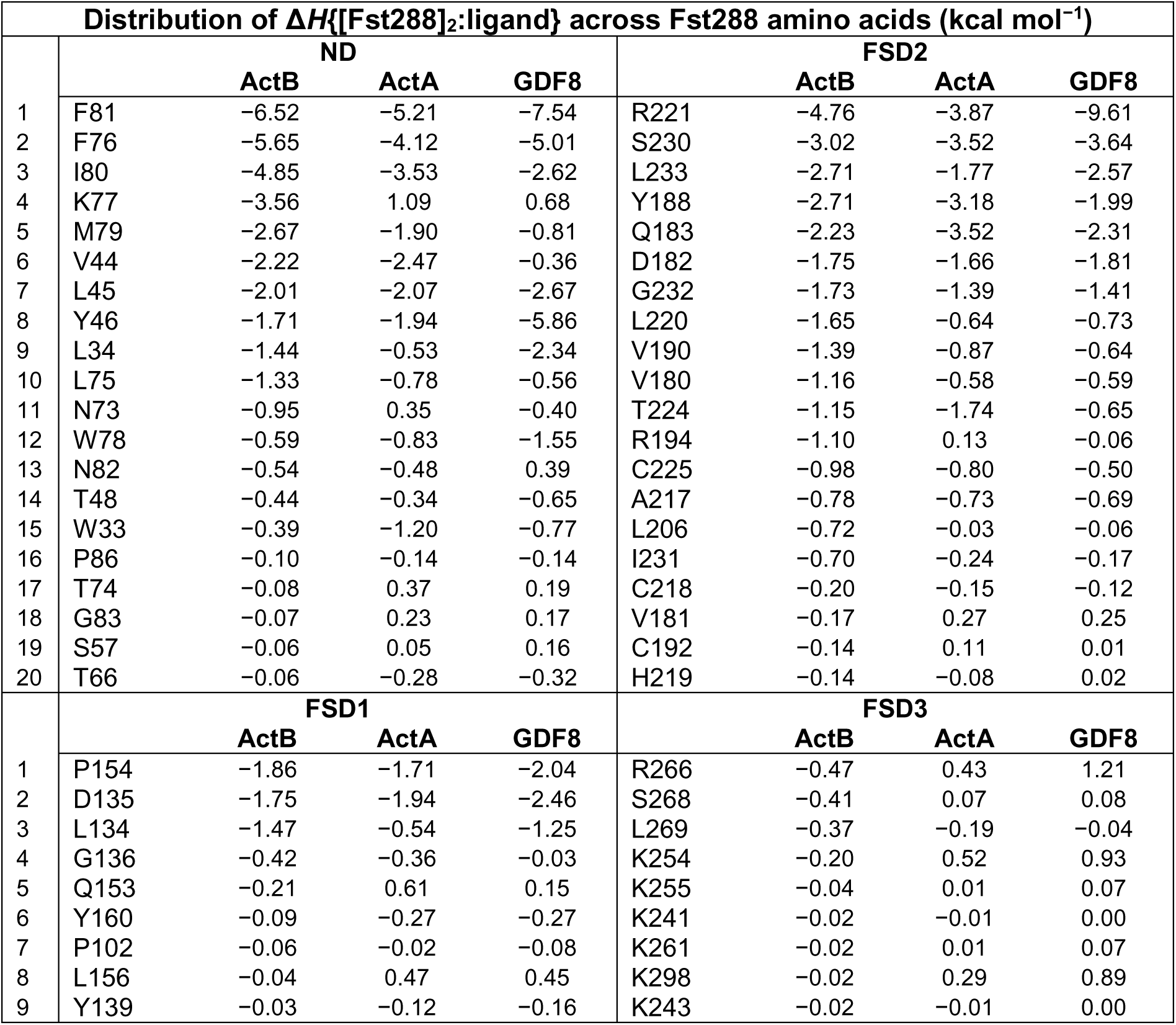
Distribution of calculated MM-GBSA enthalpy between the two Fst288 molecules and ligands across Fst288 amino acids, grouped by individual domains (in kcal mol⁻¹). Only the top 20 or 9 most contributing residues to ActB binding are shown, along with their corresponding values in the ActA and GDF8 complexes. Values represent averages of both Fst288 from five MD replicas.

**Table S4.**
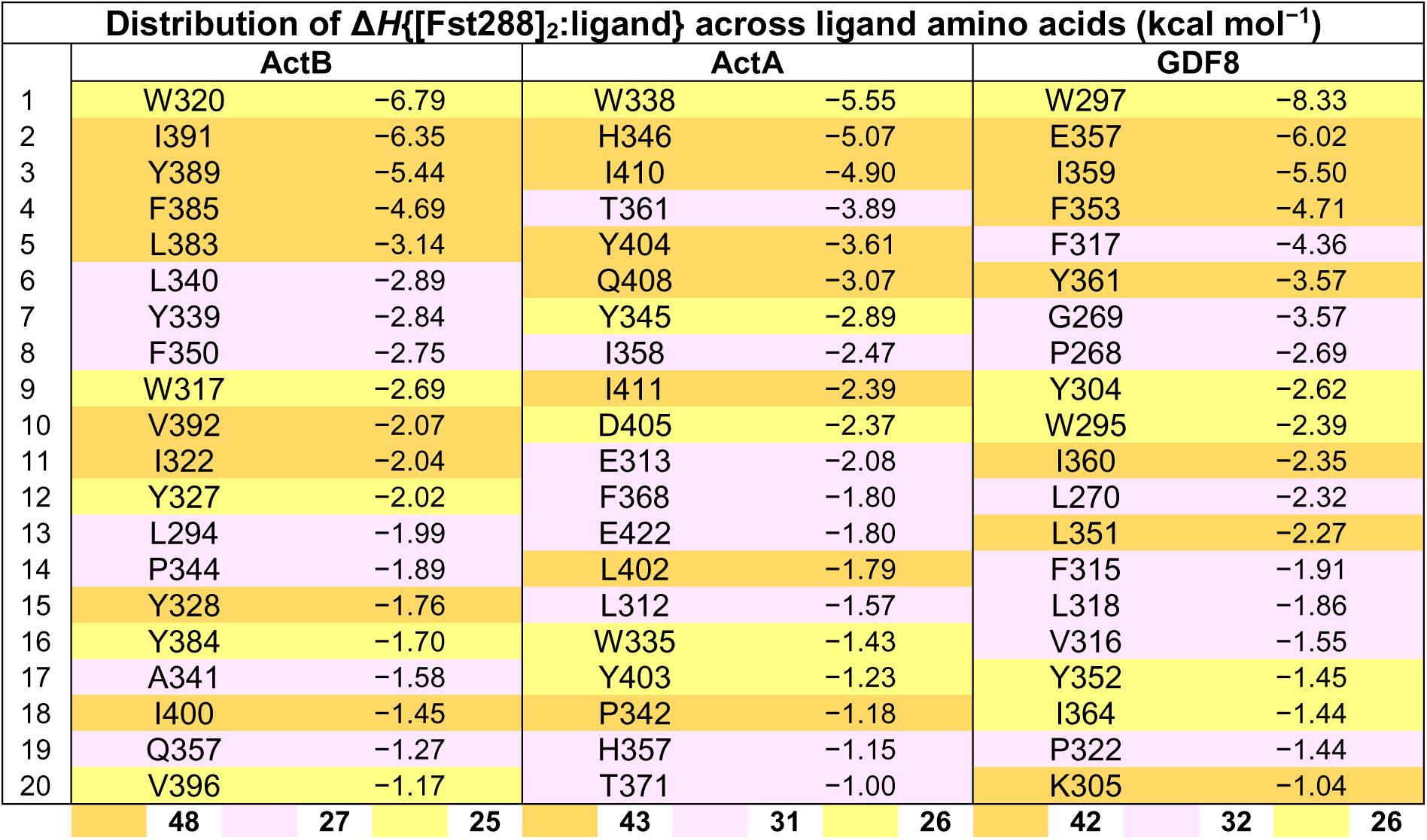
Distribution of calculated MM-GBSA enthalpy between the two Fst288 molecules and ligands across ligand amino acids (in kcal mol⁻¹). Only the top 20 most contributing residues are shown. Values represent averages of both ligand chains from five MD replicas. Residues in the type II epitope are shaded in orange, residues in the type I site from chain_1_ are shaded in yellow, and those in the type I site from chain_2_ are shaded in purple. The bottom row shows the percentage of the energy contribution to the sum of the 20 displayed amino acids from the orange, yellow, and purple regions.

| Distribution of $\Delta H\{[Fst288]_2:ligand\}$ across ligand amino acids (kcal mol<sup>-1</sup>) | | | | | | | | | |
| --- | --- | --- | --- | --- | --- | --- | --- | --- | --- |
|  | ActB |  |  | ActA |  |  | GDF8 |  |  |
| 1 | W320 | -6.79 |  | W338 | -5.55 |  | W297 | -8.33 |  |
| 2 | I391 | -6.35 |  | H346 | -5.07 |  | E357 | -6.02 |  |
| 3 | Y389 | -5.44 |  | I410 | -4.90 |  | I359 | -5.50 |  |
| 4 | F385 | -4.69 |  | T361 | -3.89 |  | F353 | -4.71 |  |
| 5 | L383 | -3.14 |  | Y404 | -3.61 |  | F317 | -4.36 |  |
| 6 | L340 | -2.89 |  | Q408 | -3.07 |  | Y361 | -3.57 |  |
| 7 | Y339 | -2.84 |  | Y345 | -2.89 |  | G269 | -3.57 |  |
| 8 | F350 | -2.75 |  | I358 | -2.47 |  | P268 | -2.69 |  |
| 9 | W317 | -2.69 |  | I411 | -2.39 |  | Y304 | -2.62 |  |
| 10 | V392 | -2.07 |  | D405 | -2.37 |  | W295 | -2.39 |  |
| 11 | I322 | -2.04 |  | E313 | -2.08 |  | I360 | -2.35 |  |
| 12 | Y327 | -2.02 |  | F368 | -1.80 |  | L270 | -2.32 |  |
| 13 | L294 | -1.99 |  | E422 | -1.80 |  | L351 | -2.27 |  |
| 14 | P344 | -1.89 |  | L402 | -1.79 |  | F315 | -1.91 |  |
| 15 | Y328 | -1.76 |  | L312 | -1.57 |  | L318 | -1.86 |  |
| 16 | Y384 | -1.70 |  | W335 | -1.43 |  | V316 | -1.55 |  |
| 17 | A341 | -1.58 |  | Y403 | -1.23 |  | Y352 | -1.45 |  |
| 18 | I400 | -1.45 |  | P342 | -1.18 |  | I364 | -1.44 |  |
| 19 | Q357 | -1.27 |  | H357 | -1.15 |  | P322 | -1.44 |  |
| 20 | V396 | -1.17 |  | T371 | -1.00 |  | K305 | -1.04 |  |
|  | 48 | 27 | 25 | 43 | 31 | 26 | 42 | 32 | 26 |

## References

1. Hinck, A. P., Mueller, T. D., and Springer, T. A. (2016) Structural Biology and Evolution of the TGF-β Family. Cold Spring Harb Perspect Biol. 8, a022103

2. Hashimoto, O., Kawasaki, N., Tsuchida, K., Shimasaki, S., Hayakawa, T., and Sugino, H. (2000) Difference between follistatin isoforms in the inhibition of activin signalling: Cellular Signalling. 12, 565–571

3. Sidis, Y., Mukherjee, A., Keutmann, H., Delbaere, A., Sadatsuki, M., and Schneyer, A. (2006) Biological Activity of Follistatin Isoforms and Follistatin-Like-3 Is Dependent on Differential Cell Surface Binding and Specificity for Activin, Myostatin, and Bone Morphogenetic Proteins. Endocrinology. 147, 3586–3597

4. Matzuk, M. M., Lu, N., Vogel, H., Sellheyer, K., Roop, D. R., and Bradley, A. (1995) Multiple defects and perinatal death in mice deficient in follistatin. Nature. 374, 360–363

5. Cash, J. N., Angerman, E. B., Kattamuri, C., Nolan, K., Zhao, H., Sidis, Y., Keutmann, H. T., and Thompson, T. B. (2012) Structure of Myostatin·Follistatin-like 3. J Biol Chem. 287, 1043–1053

6. Schneyer, A., Schoen, A., Quigg, A., and Sidis, Y. (2003) Differential binding and neutralization of activins A and B by follistatin and follistatin like-3 (FSTL-3/FSRP/FLRG). Endocrinology. 144, 1671–1674

7. Chang, C. (2016) Agonists and Antagonists of TGF-β Family Ligands. Cold Spring Harb Perspect Biol. 8, a021923

8. Thompson, T. B., Lerch, T. F., Cook, R. W., Woodruff, T. K., and Jardetzky, T. S. (2005) The Structure of the Follistatin:Activin Complex Reveals Antagonism of Both Type I and Type II Receptor Binding. Developmental Cell. 9, 535–543

9. Lerch, T. F., Shimasaki, S., Woodruff, T. K., and Jardetzky, T. S. (2007) Structural and Biophysical Coupling of Heparin and Activin Binding to Follistatin Isoform Functions*. Journal of Biological Chemistry. 282, 15930–15939

10. Stamler, R., Keutmann, H. T., Sidis, Y., Kattamuri, C., Schneyer, A., and Thompson, T. B. (2008) The Structure of FSTL3·Activin A Complex: DIFFERENTIAL BINDING OF N-TERMINAL DOMAINS INFLUENCES FOLLISTATIN-TYPE ANTAGONIST SPECIFICITY*. Journal of Biological Chemistry. 283, 32831–32838

11. Cash, J. N., Rejon, C. A., McPherron, A. C., Bernard, D. J., and Thompson, T. B. (2009) The structure of myostatin:follistatin 288: insights into receptor utilization and heparin binding. The EMBO Journal. 28, 2662–2676

12. Walker, R. G., Czepnik, M., Goebel, E. J., McCoy, J. C., Vujic, A., Cho, M., Oh, J., Aykul, S., Walton, K. L., Schang, G., Bernard, D. J., Hinck, A. P., Harrison, C. A., Martinez-Hackert, E., Wagers, A. J., Lee, R. T., and Thompson, T. B. (2017) Structural basis for potency differences between GDF8 and GDF11. BMC Biology. 15, 19

13. Keutmann, H. T., Schneyer, A. L., and Sidis, Y. (2004) The Role of Follistatin Domains in Follistatin Biological Action. Molecular Endocrinology. 18, 228–240

14. Cash, J. N., Angerman, E. B., Keutmann, H. T., and Thompson, T. B. (2012) Characterization of Follistatin-Type Domains and Their Contribution to Myostatin and Activin A Antagonism. Molecular Endocrinology. 26, 1167–1178

15. Harrington, A. E., Morris-Triggs, S. A., Ruotolo, B. T., Robinson, C. V., Ohnuma, S., and Hyvönen, M. (2006) Structural basis for the inhibition of activin signalling by follistatin. The EMBO Journal. 25, 1035–1045

16. Schneyer, A. L., Sidis, Y., Gulati, A., Sun, J. L., Keutmann, H., and Krasney, P. A. (2008) Differential antagonism of activin, myostatin and growth and differentiation factor 11 by wild-type and mutant follistatin. Endocrinology. 149, 4589–4595

17. Harrison, C. A., Chan, K. L., and Robertson, D. M. (2006) Activin-A binds follistatin and type II receptors through overlapping binding sites: generation of mutants with isolated binding activities. Endocrinology. 147, 2744–2753

18. Sidis, Y., Schneyer, A. L., Sluss, P. M., Johnson, L. N., and Keutmann, H. T. (2001) Follistatin: Essential Role for the N-terminal Domain in Activin Binding and Neutralization*. Journal of Biological Chemistry. 276, 17718–17726

19. Nakatani, M., Takehara, Y., Sugino, H., Matsumoto, M., Hashimoto, O., Hasegawa, Y., Murakami, T., Uezumi, A., Takeda, S., Noji, S., Sunada, Y., and Tsuchida, K. (2008) Transgenic expression of a myostatin inhibitor derived from follistatin increases skeletal muscle mass and ameliorates dystrophic pathology in mdx mice. FASEB J. 22, 477–487

20. Sidis, Y., Schneyer, A. L., Sluss, P. M., Johnson, L. N., and Keutmann, H. T. (2001) Follistatin: Essential Role for the N-terminal Domain in Activin Binding and Neutralization*. Journal of Biological Chemistry. 276, 17718–17726

21. Thompson, T. B., Woodruff, T. K., and Jardetzky, T. S. (2003) Structures of an ActRIIB:activin A complex reveal a novel binding mode for TGF-β ligand:receptor interactions. The EMBO Journal. 22, 1555–1566

22. Wang, X., Fischer, G., and Hyvönen, M. (2016) Structure and activation of pro-activin A. Nat Commun. 7, 12052

23. Chu, K.-Y., Malik, A., Thamilselvan, V., and Martinez-Hackert, E. (2022) Type II BMP and activin receptors BMPR2 and ACVR2A share a conserved mode of growth factor recognition. Journal of Biological Chemistry. 10.1016/j.jbc.2022.102076

24. Cotton, T. R., Fischer, G., Wang, X., McCoy, J. C., Czepnik, M., Thompson, T. B., and Hyvönen, M. (2018) Structure of the human myostatin precursor and determinants of growth factor latency. The EMBO Journal. 37, 367–383

25. Padyana, A. K., Vaidialingam, B., Hayes, D. B., Gupta, P., Franti, M., and Farrow, N. A. (2016) Crystal structure of human GDF11. Acta Cryst F. 72, 160–164

26. Goebel, E. J., Kattamuri, C., Gipson, G. R., Krishnan, L., Chavez, M., Czepnik, M., Maguire, M. C., Grenha, R., Håkansson, M., Logan, D. T., Grinberg, A. V., Sako, D., Castonguay, R., Kumar, R., and Thompson, T. B. (2022) Structures of activin ligand traps using natural sets of type I and type II TGFβ receptors. iScience. 10.1016/j.isci.2021.103590

27. Goebel, E. J., Corpina, R. A., Hinck, C. S., Czepnik, M., Castonguay, R., Grenha, R., Boisvert, A., Miklossy, G., Fullerton, P. T., Matzuk, M. M., Idone, V. J., Economides, A. N., Kumar, R., Hinck, A. P., and Thompson, T. B. (2019) Structural characterization of an activin class ternary receptor complex reveals a third paradigm for receptor specificity. Proceedings of the National Academy of Sciences. 116, 15505–15513

28. Namwanje, M., and Brown, C. W. (2016) Activins and Inhibins: Roles in Development, Physiology, and Disease. Cold Spring Harb Perspect Biol. 8, a021881

29. Wijayarathna, R., and de Kretser, D. M. (2016) Activins in reproductive biology and beyond. Hum Reprod Update. 22, 342–357

30. Fang, D. Y. P., Lu, B., Hayward, S., de Kretser, D. M., Cowan, P. J., and Dwyer, K. M. (2016) The Role of Activin A and B and the Benefit of Follistatin Treatment in Renal Ischemia-Reperfusion Injury in Mice. Transplant Direct. 2, e87

31. Krepinsky, J. C. (2022) Activin B, a new player in kidney fibrosis?†. J Pathol. 256, 363–365

32. Nagayama, I., Takei, Y., Takahashi, S., Okada, M., and Maeshima, A. (2025) The activin-follistatin system: Key regulator of kidney development, regeneration, inflammation, and fibrosis. Cytokine Growth Factor Rev. 81, 1–8

33. Thompson, T. B., Cook, R. W., Chapman, S. C., Jardetzky, T. S., and Woodruff, T. K. (2004) Beta A versus beta B: is it merely a matter of expression? Mol Cell Endocrinol. 225, 9–17

34. Nolan, K., Kattamuri, C., Rankin, S. A., Read, R. J., Zorn, A. M., and Thompson, T. B. (2016) Structure of Gremlin-2 in Complex with GDF5 Gives Insight into DAN-Family-Mediated BMP Antagonism. Cell Reports. 16, 2077–2086

35. Mast, E. M., Leach, E. A. E., and Thompson, T. B. (2024) Characterization of erythroferrone oligomerization and its impact on BMP antagonism. J Biol Chem. 300, 105452

36. Yariv, B., Yariv, E., Kessel, A., Masrati, G., Chorin, A. B., Martz, E., Mayrose, I., Pupko, T., and Ben-Tal, N. (2023) Using evolutionary data to make sense of macromolecules with a “face-lifted” ConSurf. Protein Sci. 32, e4582

37. Walker, R. G., Kattamuri, C., Goebel, E. J., Zhang, F., Hammel, M., Tainer, J. A., Linhardt, R. J., and Thompson, T. B. (2021) Heparin-mediated dimerization of follistatin. Exp Biol Med (Maywood). 246, 467–482

38. Zhang, F., Beaudet, J. M., Luedeke, D. M., Walker, R. G., Thompson, T. B., and Linhardt, R. J. (2012) Analysis of the Interaction between Heparin and Follistatin and Heparin and Follistatin–Ligand Complexes Using Surface Plasmon Resonance. Biochemistry. 51, 6797–6803

39. Ueno, N., Ling, N., Ying, S. Y., Esch, F., Shimasaki, S., and Guillemin, R. (1987) Isolation and partial characterization of follistatin: a single-chain Mr 35,000 monomeric protein that inhibits the release of follicle-stimulating hormone. Proc Natl Acad Sci U S A. 84, 8282–8286

40. Hashimoto, O., Nakamura, T., Shoji, H., Shimasaki, S., Hayashi, Y., and Sugino, H. (1997) A novel role of follistatin, an activin-binding protein, in the inhibition of activin action in rat pituitary cells. Endocytotic degradation of activin and its acceleration by follistatin associated with cell-surface heparan sulfate. J Biol Chem. 272, 13835–13842

41. McCoy, A. J., Grosse-Kunstleve, R. W., Adams, P. D., Winn, M. D., Storoni, L. C., and Read, R. J. (2007) Phaser crystallographic software. J Appl Crystallogr. 40, 658–674

42. Agirre, J., Atanasova, M., Bagdonas, H., Ballard, C. B., Baslé, A., Beilsten-Edmands, J., Borges, R. J., Brown, D. G., Burgos-Mármol, J. J., Berrisford, J. M., Bond, P. S., Caballero, I., Catapano, L., Chojnowski, G., Cook, A. G., Cowtan, K. D., Croll, T. I., Debreczeni, J. É., Devenish, N. E., Dodson, E. J., Drevon, T. R., Emsley, P., Evans, G., Evans, P. R., Fando, M., Foadi, J., Fuentes-Montero, L., Garman, E. F., Gerstel, M., Gildea, R. J., Hatti, K., Hekkelman, M. L., Heuser, P., Hoh, S. W., Hough, M. A., Jenkins, H. T., Jiménez, E., Joosten, R. P., Keegan, R. M., Keep, N., Krissinel, E. B., Kolenko, P., Kovalevskiy, O., Lamzin, V. S., Lawson, D. M., Lebedev, A. A., Leslie, A. G. W., Lohkamp, B., Long, F., Malý, M., McCoy, A. J., McNicholas, S. J., Medina, A., Millán, C., Murray, J. W., Murshudov, G. N., Nicholls, R. A., Noble, M. E. M., Oeffner, R., Pannu, N. S., Parkhurst, J. M., Pearce, N., Pereira, J., Perrakis, A., Powell, H. R., Read, R. J., Rigden, D. J., Rochira, W., Sammito, M., Sánchez Rodríguez, F., Sheldrick, G. M., Shelley, K. L., Simkovic, F., Simpkin, A. J., Skubak, P., Sobolev, E., Steiner, R. A., Stevenson, K., Tews, I., Thomas, J. M. H., Thorn, A., Valls, J. T., Uski, V., Usón, I., Vagin, A., Velankar, S., Vollmar, M., Walden, H., Waterman, D., Wilson, K. S., Winn, M. D., Winter, G., Wojdyr, M., and Yamashita, K. (2023) The CCP4 suite: integrative software for macromolecular crystallography. Acta Cryst D. 79, 449–461

43. Abramson, J., Adler, J., Dunger, J., Evans, R., Green, T., Pritzel, A., Ronneberger, O., Willmore, L., Ballard, A. J., Bambrick, J., Bodenstein, S. W., Evans, D. A., Hung, C.-C., O’Neill, M., Reiman, D., Tunyasuvunakool, K., Wu, Z., Žemgulytė, A., Arvaniti, E., Beattie, C., Bertolli, O., Bridgland, A., Cherepanov, A., Congreve, M., Cowen-Rivers, A. I., Cowie, A., Figurnov, M., Fuchs, F. B., Gladman, H., Jain, R., Khan, Y. A., Low, C. M. R., Perlin, K., Potapenko, A., Savy, P., Singh, S., Stecula, A., Thillaisundaram, A., Tong, C., Yakneen, S., Zhong, E. D., Zielinski, M., Žídek, A., Bapst, V., Kohli, P., Jaderberg, M., Hassabis, D., and Jumper, J. M. (2024) Accurate structure prediction of biomolecular interactions with AlphaFold 3. Nature. 630, 493–500

44. Liebschner, D., Afonine, P. V., Baker, M. L., Bunkóczi, G., Chen, V. B., Croll, T. I., Hintze, B., Hung, L. W., Jain, S., McCoy, A. J., Moriarty, N. W., Oeffner, R. D., Poon, B. K., Prisant, M. G., Read, R. J., Richardson, J. S., Richardson, D. C., Sammito, M. D., Sobolev, O. V., Stockwell, D. H., Terwilliger, T. C., Urzhumtsev, A. G., Videau, L. L., Williams, C. J., and Adams, P. D. (2019) Macromolecular structure determination using X-rays, neutrons and electrons: recent developments in Phenix. Acta Crystallogr D Struct Biol. 75, 861–877

45. Emsley, P., Lohkamp, B., Scott, W. G., and Cowtan, K. (2010) Features and development of Coot. Acta Crystallogr D Biol Crystallogr. 66, 486–501

46. Šali, A., and Blundell, T. L. (1993) Comparative Protein Modelling by Satisfaction of Spatial Restraints. Journal of Molecular Biology. 234, 779–815

47. Olsson, M. H. M., Søndergaard, C. R., Rostkowski, M., and Jensen, J. H. (2011) PROPKA3: Consistent Treatment of Internal and Surface Residues in Empirical pKa Predictions. J. Chem. Theory Comput. 7, 525–537

48. Tian, C., Kasavajhala, K., Belfon, K. A. A., Raguette, L., Huang, H., Migues, A. N., Bickel, J., Wang, Y., Pincay, J., Wu, Q., and Simmerling, C. (2020) ff19SB: Amino-Acid-Specific Protein Backbone Parameters Trained against Quantum Mechanics Energy Surfaces in Solution. J. Chem. Theory Comput. 16, 528–552

49. Izadi, S., Anandakrishnan, R., and Onufriev, A. V. (2014) Building Water Models: A Different Approach. J. Phys. Chem. Lett. 5, 3863–3871

50. An overview of the Amber biomolecular simulation package - Salomon-Ferrer - 2013 - WIREs Computational Molecular Science - Wiley Online Library [online] https://wires.onlinelibrary.wiley.com/doi/10.1002/wcms.1121 (Accessed November 11, 2025)

51. Yan, Y., Tao, H., He, J., and Huang, S.-Y. (2020) The HDOCK server for integrated protein–protein docking. Nat Protoc. 15, 1829–1852

52. Roe, D. R., and Cheatham, T. E. I. (2013) PTRAJ and CPPTRAJ: Software for Processing and Analysis of Molecular Dynamics Trajectory Data. J. Chem. Theory Comput. 9, 3084–3095

53. Genheden, S., and Ryde, U. (2015) The MM/PBSA and MM/GBSA methods to estimate ligand-binding affinities. Expert Opin Drug Discov. 10, 449–461

54. Reliable In Silico Ranking of Engineered Therapeutic TCR Binding Affinities with MMPB/GBSA | Journal of Chemical Information and Modeling [online] https://pubs.acs.org/doi/10.1021/acs.jcim.1c00765 (Accessed November 11, 2025)

55. Abraham, M. J., Murtola, T., Schulz, R., Páll, S., Smith, J. C., Hess, B., and Lindahl, E. (2015) GROMACS: High performance molecular simulations through multi-level parallelism from laptops to supercomputers. SoftwareX. 1–2, 19–25

56. Hub, J. S., de Groot, B. L., and van der Spoel, D. (2010) g_wham—A Free Weighted Histogram Analysis Implementation Including Robust Error and Autocorrelation Estimates. J. Chem. Theory Comput. 6, 3713–3720

